# ENGINEERING, DECODING AND SYSTEMS-LEVEL CHARACTERIZATION OF CHIMPANZEE CYTOMEGALOVIRUS

**DOI:** 10.1101/2021.07.20.453063

**Authors:** Quang Vinh Phan, Boris Bogdanow, Emanuel Wyler, Markus Landthaler, Fan Liu, Christian Hagemeier, Lüder Wiebusch

## Abstract

The chimpanzee cytomegalovirus (CCMV) is the closest relative of human CMV (HCMV). Because of the high conservation between these two species and the ability of human cells to fully support CCMV replication, CCMV holds great potential as a model system for HCMV. To make the CCMV genome available for precise and rapid gene manipulation techniques, we captured the genomic DNA of CCMV strain Heberling as a bacterial artificial chromosome (BAC). Selected BAC clones were reconstituted to infectious viruses, growing to similar high titers as parental CCMV. DNA sequencing confirmed the integrity of our clones and led to the identification of two polymorphic loci within the CCMV genome. To re-evaluate the CCMV coding potential, we analyzed the transcriptome and proteome of infected cells and identified several novel ORFs, splice variants, and regulatory RNAs. We further characterized the dynamics of CCMV gene expression and found that viral proteins cluster into five distinct temporal classes. In addition, our datasets revealed that the host response to CCMV infection and the de-regulation of cellular pathways are in line with known hallmarks of HCMV infection. In a first functional experiment, we investigated a proposed frameshift mutation in UL128 that was suspected to restrict CCMV’s cell tropism. In fact, repair of this frameshift re-established productive CCMV infection in endothelial and epithelial cells, expanding the options of CCMV as an infection model. Thus, BAC-cloned CCMV can serve as a powerful tool for systematic approaches in comparative functional genomics, exploiting the close phylogenetic relationship between CCMV and HCMV.

## INTRODUCTION

Cytomegaloviruses (CMVs) are a group of β-herpesviruses infecting numerous primate species, including old and new world monkeys, great apes and humans (Mozzi et al., 2020). As a result of prolonged periods of coevolution, CMVs are highly adapted to their specific hosts, making cross-species infections extremely rare events (Burwitz et al., 2016; Child et al., 2021; Murthy et al., 2019). CMVs are very prevalent in their host populations, due to life-long, mostly asymptomatic infections. Medically most relevant is the human cytomegalovirus (HCMV) which causes serious complications in immunocompromised patients and neonates. Available HCMV treatment relies on inhibitors of viral DNA replication and packaging but their use is limited by side effects and the emergence of resistant virus strains. Currently, there are no approved HCMV vaccines available.

The severe consequences on health and the limitation in treatment options have motivated systematic efforts in understanding HCMV infection and replication mechanisms. Transcriptomic and proteomic studies have provided a comprehensive picture of the viral gene expression program (Gatherer et al., 2011; Gonzalez-Perez et al., 2021; Marcinowski et al., 2012; Stern-Ginossar et al., 2012; Tirosh et al., 2015; Weekes et al., 2014). This has been complemented by global assessment of virus-virus and virus-host protein interactions (Hernández Durán et al., 2019; Nobre et al., 2019). By genome-wide mutagenesis of viral DNA, a set of genes essential for HCMV replication *in vitro* has been identified, which mostly consists of “core” genes conserved throughout herpesviruses (Dunn et al., 2003; Hein and Weissman, 2021; Yu et al., 2003). In addition, HCMV encodes an arsenal of accessory, less-conserved factors required for immune evasion (Dell’Oste et al., 2020; Patro, 2019), regulation of host metabolism (Rodríguez-Sánchez and Munger, 2019), cell cycle control (Spector, 2015) and anti-apoptosis (Brune and Andoniou, 2017).

For the functional analysis of potential antiviral targets by reverse genetics, CMV genomes cloned as bacterial artificial chromosomes (BACs) serve as powerful tools (Brune et al., 2000; Tischer and Kaufer, 2012). First, BAC-captured CMV genomes are easily propagated and stably maintained as single clones in *Escherichia coli* (*E*.*coli*). Second, and more important, the large and complex CMV genomes become available for prokaryotic recombination techniques, allowing for efficient and reliable gene targeting. In addition to multiple HCMV strains cloned as BACs (Wilkinson et al., 2015), animal CMV BACs are utilized as cloning platforms for *in vitro* and *in vivo* models and thereby provide the framework for a profound understanding of CMV infection and pathogenesis (Roark et al., 2020; Taher et al., 2020). In particular, the rhesus CMV (RhCMV) BAC platform has become a popular model system for HCMV due to many shared features in infection, replication and immune control (Itell et al., 2017; Powers and Früh, 2008).

RhCMV proteins show only a moderate level of conservation towards HCMV homologs at the amino acid level (Barry and William Chang, 2007). Furthermore, in two genomic regions, comprising the RL11 gene family and the ULb’ locus (Umashankar et al., 2011), the genetic content of RhCMV diverges substantially from HCMV (Malouli et al., 2012). This degree of evolutionary divergence can set a limitation on the ability to assess HCMV gene functions by studying RhCMV homologs. By comparison, chimpanzee CMV (CCMV) shares a much higher level of similarity in both genome organization and protein content with HCMV (Barry and William Chang, 2007; Malouli et al., 2012; Mozzi et al., 2020). The close phylogenetic relationship between HCMV and CCMV (Alcendor et al., 2009; Leendertz et al., 2009) is reflected by the fact that human cells are fully permissive for CCMV and vice versa (Davison et al., 2003; Perot et al., 1992). Accordingly, whole genome sequencing data of the CCMV Heberling strain were utilized to re-evaluate the HCMV coding potential (Davison et al., 2003) and to investigate the adaptation of HCMV to its human host (Mozzi et al., 2020).

We envisioned CCMV to be an eminent model system that operates at the interface between animal and human CMV species and therefore BAC-cloned the CCMV Heberling genome. DNA sequencing confirmed the integrity of our clones and led to the identification of two polymorphic loci within the CCMV genome. To re-evaluate the CCMV coding potential, we analyzed the transcriptome and proteome of infected cells and identified several novel translated open reading frames (ORFs), splice variants, and regulatory RNAs. When characterizing the dynamics of CCMV gene expression, the host response to CCMV infection and the de-regulation of cellular pathways, we found many parallels with known hallmarks of HCMV infection. Our system-level characterization of a clonal CCMV strain thus reinforces the potential of CCMV as a model system for HCMV infection.

## RESULTS

### BAC cloning of CCMV Heberling strain

To make CCMV accessible for efficient genetic analyses, we set out to clone the CCMV strain Heberling genome as a bacterial artificial chromosome (BAC). Due to the high collinearity between HCMV and CCMV, we chose a cloning strategy similar to the approach taken for the generation of HCMV-TB40-BAC4 (Christian Sinzger et al., 2008). To this end, the BAC donor plasmid pEB1097 (Borst et al., 1999) was modified in a way that integration of the BAC cassette via homologous recombination replaces the US2 to US6 region of CCMV (Fig. 1A). After enrichment of recombined CCMV genomes by guanine phosphoribosyltransferase selection, circular viral DNA was extracted and transformed into E. coli. To check the overall integrity of the CCMV genome, randomly selected BAC clones were analyzed by AgeI restriction digest. Five clones closely matched the *in silico* predicted restriction pattern (Fig. 1B) and were further analyzed for infectivity and virus growth. Upon re-transfection into primary human fibroblasts, all clones reconstituted to replication-competent viruses. Although their growth kinetics were similar to the parental Heberling virus, the BAC-derived viruses reached a lower maximum titer (Fig. 1C).

**FIG 1.**
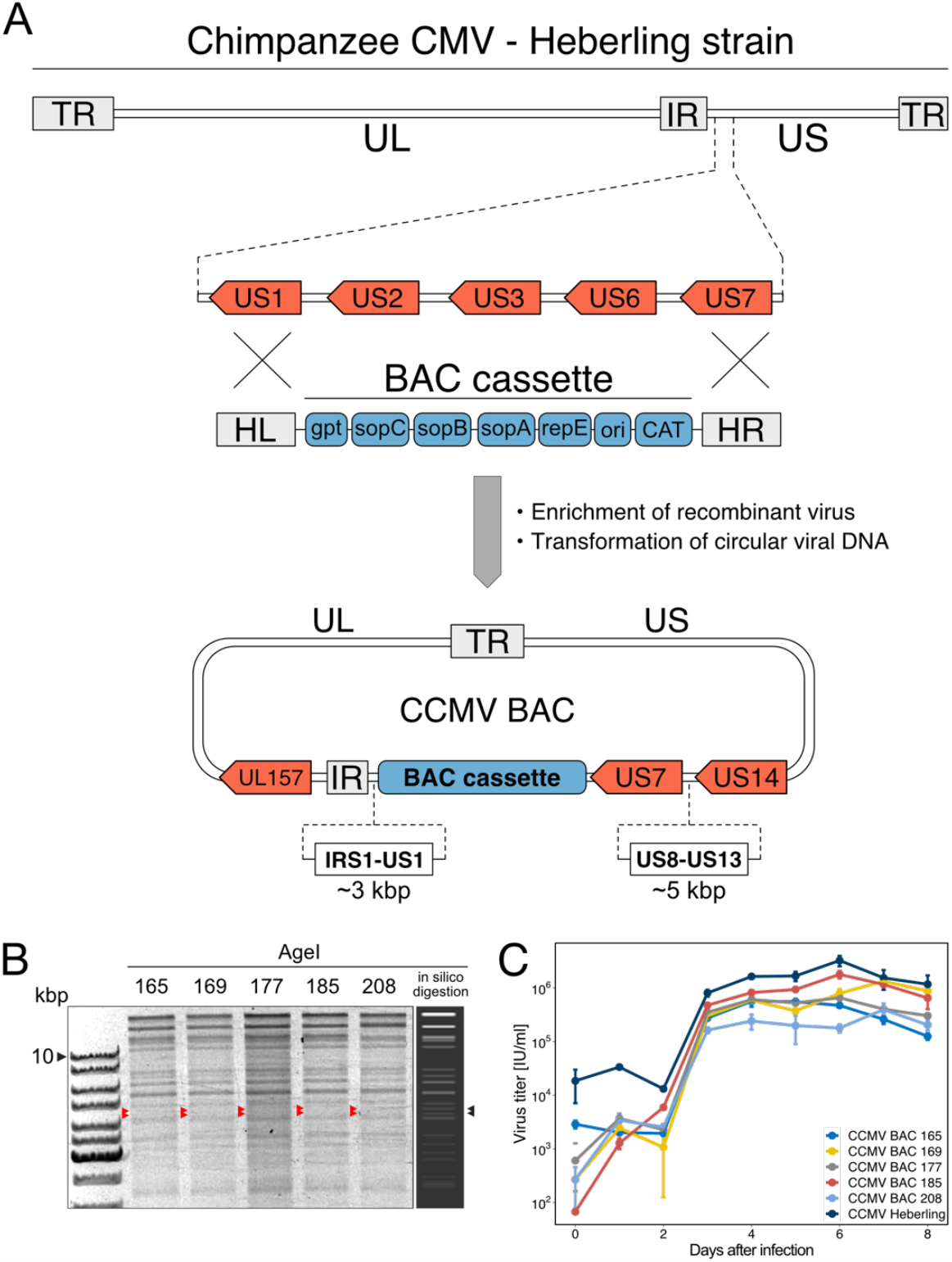
BAC-cloning of CCMV strain Heberling. (A) Targeted integration of the BAC cassette into the CCMV Heberling genome was mediated by homology arms left (HL) and right (HR), replacing the region between US1 and US7. The BAC cassette contains a replication origin (ori), a centromeric region (sopC), genes encoding replication (repE) and partitioning factors (sopA, sopB), as well as resistance genes for eukaryotic (xanthine phosphoribosyltransferase - gpt) and prokaryotic selection (chloramphenicol acetyltransferase - CAT). TR - terminal repeats; UL - unique long region; IR - internal repeats; US - unique short region. (B) Five BAC clones showed an AgeI restriction pattern matching the predicted *in silico* digest. Arrowheads indicate bands that were missing due to an unintended deletion of IRS1 to US1 and US8 to US13 regions. (C) All selected CCMV BAC clones were infectious upon transfection into fibroblast but differ in their titer output performance. Means (center of the error bar) and standard errors of the mean of n=3 are depicted.

Compared to the *in silico* prediction, two AgeI restriction fragments of 4272 and 4560 bp length were missing in the analytical digests of our CCMV-BAC clones (see arrowheads in Fig. 1B). As these fragments originate from the US region of the CCMV genome, we had to consider the possibility that the targeted integration of the BAC cassette was not as accurate as expected. We therefore performed polymerase chain reaction (PCR) assays and Sanger sequencing to validate the anticipated recombination event. In all clones we found an approximately 3.6 kbp genomic deletion adjacent to the 5’-end of the BAC cassette (Fig. S5A) and an additional 5.2 kbp deletion in the vicinity of the 3’-end (Fig. S5B). This finding was reminiscent of the accidental loss of the IRS1-US1 region that has occurred during the generation of TB40-BAC4 (Sampaio et al., 2017). To characterize the genomes of the selected CCMV BAC clones in more detail, we subjected them to whole genome sequencing. The alignment of sequencing reads to the reference genome confirmed the unintended deletion events of the IRS1-US1 and US8-US13 regions. Importantly, the overall CCMV genome integrity was maintained as no other major deletion or insertion (indels) events were detected (Fig. S5C).

The whole genome sequencing data allowed an in-depth genotypic analysis of the CCMV-BAC clones. In comparison to the published CCMV Heberling sequence (Davison et al., 2003), we observed an accumulation of single nucleotide variations (SNV) within the UL48-55 and the US26-28 regions (Fig. 2A, S6, Table S1). To answer whether these mutations arose during the BAC cloning procedure or were already present prior to our cloning efforts, we included DNA sequencing data of the parental Heberling stock in our analysis. As CCMV Heberling had never been plaque purified (Davison et al., 2003), it presented as a mixture of different genotypes. The nucleotide variations between these genotypes clustered at the same genomic positions as in our BAC clones (Fig. 2A, Data Set S1). In total, at least 62% (419/681) of SNV were carried over from the viral stock to our clones, leaving only a minor fraction of mutations that were likely to be created *de novo* during BAC generation.

**FIG 2.**
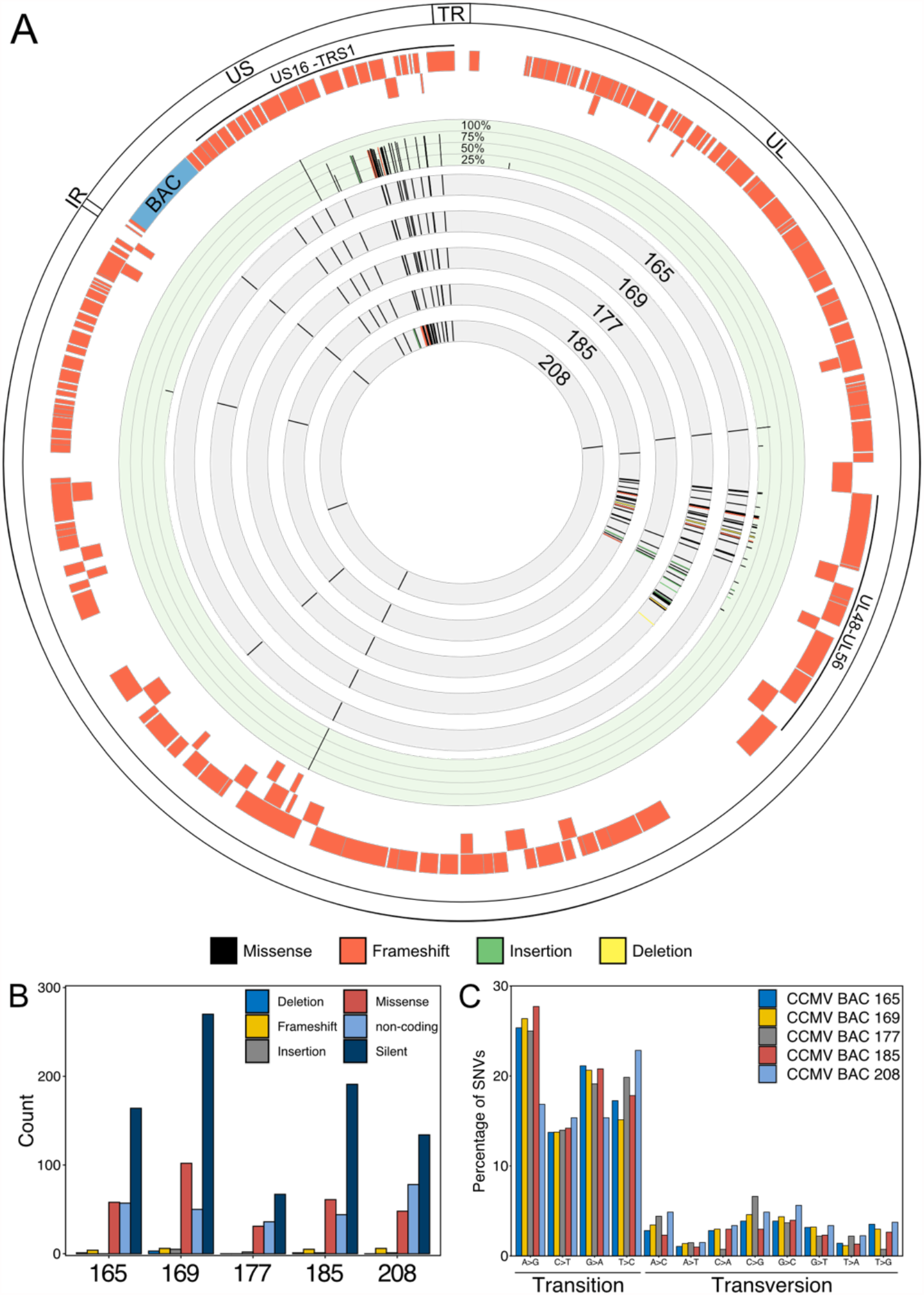
Genotypic analysis of CCMV clones. (A) Whole genome sequencing revealed variations in genomic sequences of selected CCMV BAC clones and the parental CCMV Heberling. Positions of missense, frameshift, deletion and insertion mutations for BAC clones are plotted in the grey tracks of the Circos plot. Intra-strain variations of the CCMV Heberling isolate are plotted in the green track; the frequency of each variation is indicated. Outer tracks: CCMV ORFs (red) and genomic regions. (B) The individual BAC clones were compared with the CCMV Heberling reference sequence for the absolute counts of single nucleotide variations (SNVs). CCMV BAC-177 showed the lowest number of SNVs and contained no frameshift mutations. (C) Similarly, the BAC clones were analyzed for the spectrum of transition and transversion mutations.

Silent mutations contributed the major proportion of SNVs in the analyzed CCMV sequences (Fig. 2B, Fig. S6). This was in line with the observation that base transitions are more prevalent than base transversions (Fig. 2C). The number of missense mutations ranged from 31 (BAC-177) to 102 (BAC-169) and 4 to 6 frameshift mutations were found in all BAC sequences except BAC-177. Taken together, BAC clone 177 was the most accurate representation of the originally published CCMV Heberling genome and we therefore chose this clone as a basis for further modifications.

### UL128 repair expands the cell tropism of CCMV

Fibroblasts are the preferred cell type for CMV isolation and propagation. However, extensive passage of HCMV in fibroblasts frequently results in disruptive mutations in genes of the pentameric complex (UL128, UL130, UL131A), rendering fibroblast-adapted virus strains unable to enter endothelial, epithelial and myeloid cells (C. Sinzger et al., 2008; Wilkinson et al., 2015). In consistence with this, a frameshift mutation within the first exon of the CCMV UL128 gene has been proposed for CCMV Heberling (Akter et al., 2003; Davison et al., 2003). The proposed mutation was carried over to our BAC clones, but no further disruptive mutations were detected in the members of the pentameric complex. We suspected that the frameshift mutation restricts the cell tropism of CCMV and aimed to restore UL128 function by BAC mutagenesis. After removing the cytosine insertion at position 111 of the UL128 coding sequence (Fig. 3A), we observed a substantial reduction in cell-free titer upon virus reconstitution in human fibroblasts. This was consistent with previous observations on HCMV-UL128 (Murrell et al., 2017). We then used low MOI conditions to infect human fibroblasts, epithelial and endothelial cells. In fibroblasts, the onset of viral gene expression occurred independent of the UL128 status. In contrast, only the UL128-repaired virus was capable of inducing IE gene expression and establishing virus growth in endothelial and epithelial cells (Fig. 3B-C). This data confirmed the UL128 defect in the original CCMV Heberling isolate. By reinstating the endothelial cell tropism of CCMV, we expanded the options of our CCMV-BAC for future experimental applications and provided a proof of concept that it is suitable for functional genomics approaches.

**FIG 3.**
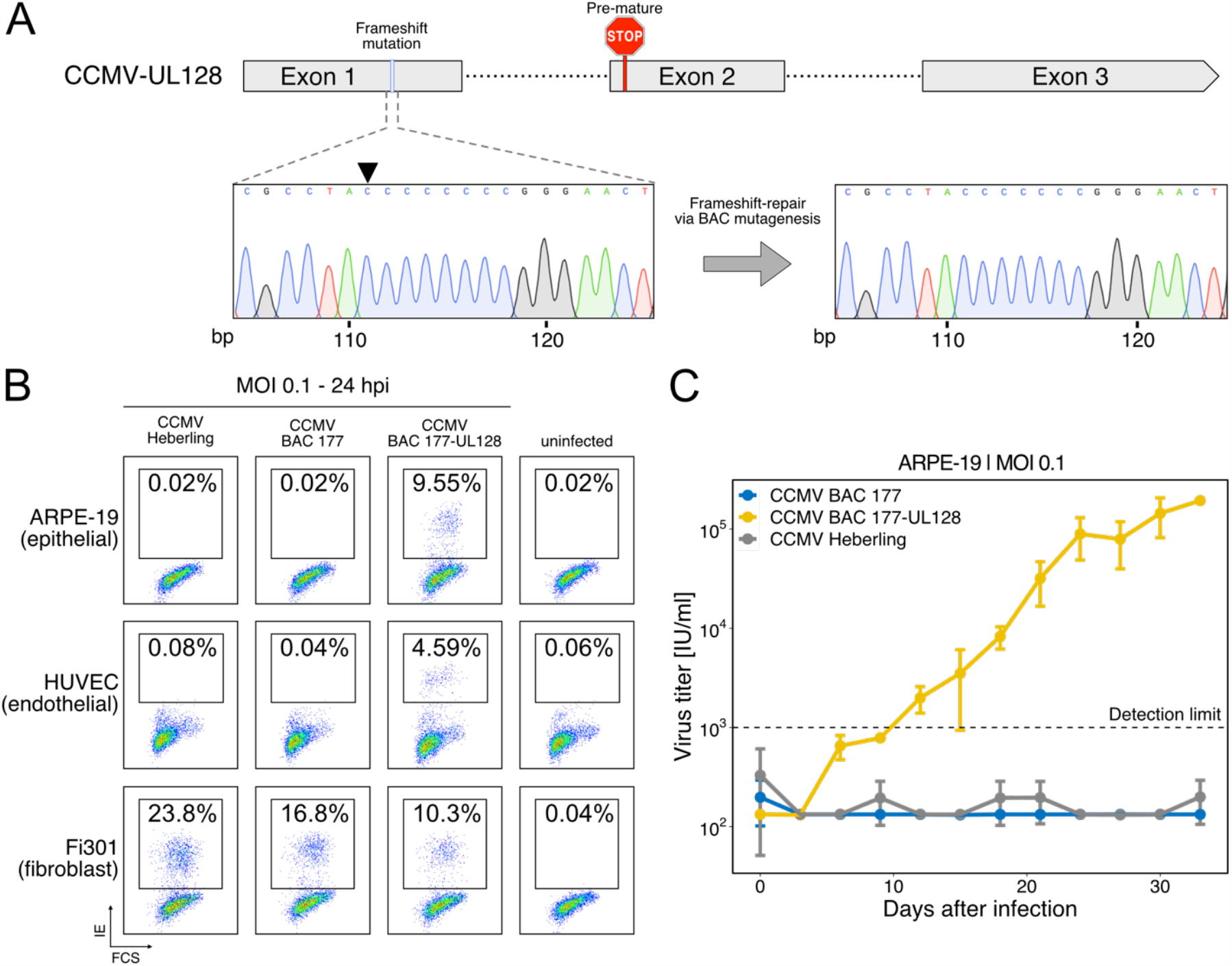
UL128 repair rescues CCMV’s tropism for epithelial and endothelial cells. (A) In CCMV Heberling, a frameshift mutation in the first exon of UL128 leads to a premature stop codon in exon 2. Sanger sequencing chromatograms of BAC-177 and the UL128-repaired BAC-177 document the successful removal of the frameshift by BAC mutagenesis. (B, C) The indicated cell types were infected with viruses derived either from the Heberling isolate, BAC-177 or the UL128-repaired BAC-177, using a multiplicity of infection (MOI) of 0.1 infectious units per cell. At 24 h post infection, the cells were analyzed for IE1/2 gene expression by flow cytometry (B). Virus growth in epithelial cells was monitored over a 33-day period. Means (center of the error bar) and standard errors of the mean of n=3 are depicted (C).

### Restoration of the US region

As mentioned above, all CCMV BAC clones displayed major deletions next to the BAC cassette integration site. Although the affected US1-US13 gene region is not essential for cytomegalovirus replication in vitro (Kollert-Jöns et al., 1991), it plays an important role in immune modulation (Berry et al., 2020). To make CCMV-BAC clone 177 useful for immunological research, we decided to restore the US region and make the BAC cassette excisable by Cre recombination (Fig. 4A). The latter step was necessary to ensure that the BAC size does not exceed the packaging capacity of the viral capsid. While the seamless restoration of the US2-US13 region was readily accomplished by BAC mutagenesis, we did not succeed in re-introducing US1 and IRS1. Possibly, the close proximity to the internal repeat region resulted in promiscuous, non-specific recombination events. The BAC cassette was flanked with loxP sites for Cre-mediated excision which only left a single loxP site between US6 and US7 behind (Fig. 4B). We verified the integrity of our improved CCMV-BAC construct, named CCMV-Phan9-BAC, via whole genome sequencing and detected 20 minor deviations from the adjusted CCMV BAC 177 reference sequence in the modified US/TR region (Fig. 4A). None of these changes had an impact on protein coding sequences and the CCMV-Phan9-BAC grew to similar titers as the parental BAC-177 virus (Fig. 4C). In the following, the CCMV-Phan9-BAC was utilized for a comprehensive characterization of the CCMV gene expression program.

**FIG 4.**
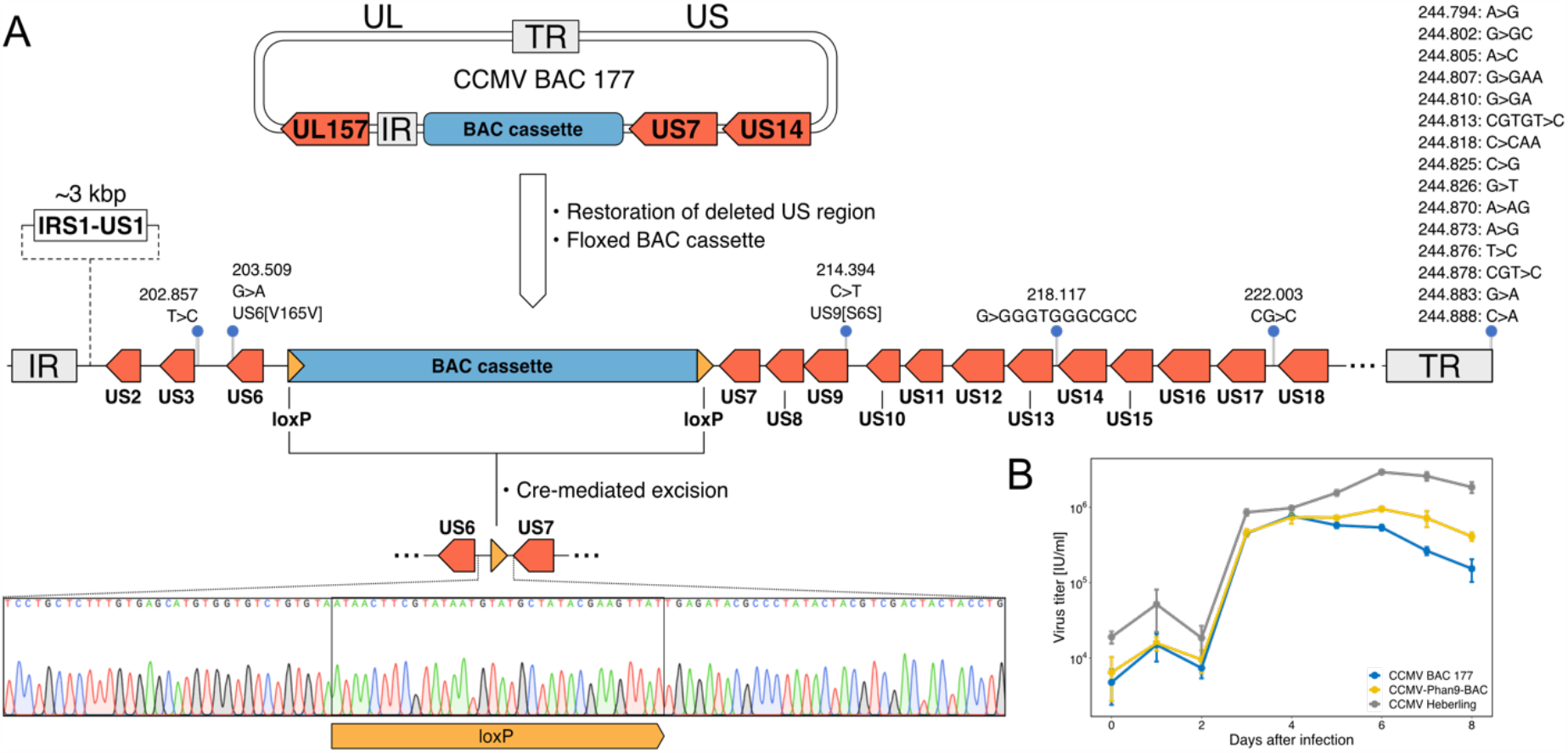
Restoration of deleted US regions and floxing of BAC cassette. (A) Schematic overview of US2-US6 and US8-US13 regions that were restored by BAC mutagenesis in CCMV BAC-177. Region IRS1 to US1 was not reinstated. LoxP sites were inserted at both ends of the BAC cassette to remove prokaryotic sequences by Cre recombinase expression during virus reconstitution. The final CCMV BAC construct was named CCMV-Phan9-BAC. Whole genome sequencing verified the integrity of CCMV Phan9-BAC. Coverage of DNA sequencing and positions of minor sequence variations in the US and TR regions are plotted. (B) Sanger sequencing of DNA from reconstituted virus confirms the expected Cre-mediated recombination outcome at the US6-US7 region. (C) One-step growth curves of CCMV BAC-177, CCMV Phan9-BAC and CCMV Heberling were analyzed in human fibroblasts. Means (center of the error bar) and standard errors of the mean of n=3 are depicted.

### Evaluation of the CCMV coding capacity

Since the initial sequencing of the CCMV genome (Davison et al., 2003), the understanding of the genetic content of other CMV genomes has been dramatically improved (Malouli et al., 2012; Stern-Ginossar et al., 2012). In contrast, CCMV gene expression has never been analyzed systematically. In order to get a deeper understanding of CCMV infection and the evolutionary relationship of CCMV and HCMV, we took our cloned virus and analyzed the CCMV coding potential by state of the art transcriptomics and proteomics. The RNA and protein material we used for this purpose was isolated from productively infected cells at five different time points, covering the whole CCMV replication cycle. First, we created a preliminary annotation file of potential novel open reading frames (ORFs) by the following stepwise approach (Fig. 5A): (i) The CCMV genome was scanned by tBLASTn (Gertz et al., 2006) for homologs of previously identified non-canonical ORFs of HCMV (Stern-Ginossar et al., 2012); out of 54 input ORFs, we identified 29 CCMV homologs. (ii) Our RNA sequencing reads were aligned to the CCMV reference genome; this revealed the presence of 45 splice junctions. By examining these splice junctions for nearby start and stop codons, 28 new ORFs were annotated. (iii) Computational prediction of ORFs was performed using the smORF software (Durrant and Bhatt, 2021) and a 6-frame annotation; this added 22 additional ORFs to our list. In total, we found 79 potential ORFs (Table S2) that were not annotated before (Davison et al., 2003). To validate their translation to proteins, we scanned our proteomic data set for matching peptides. In several instances, only peptides were detected that are shared with canonical ORFs and therefore could not be assigned unambiguously. We ended up with 14 newly annotated ORFs, to which at least two unique peptides could be assigned, proving the existence of translation products (Fig. 5A, Table S2).

**FIG 5.**
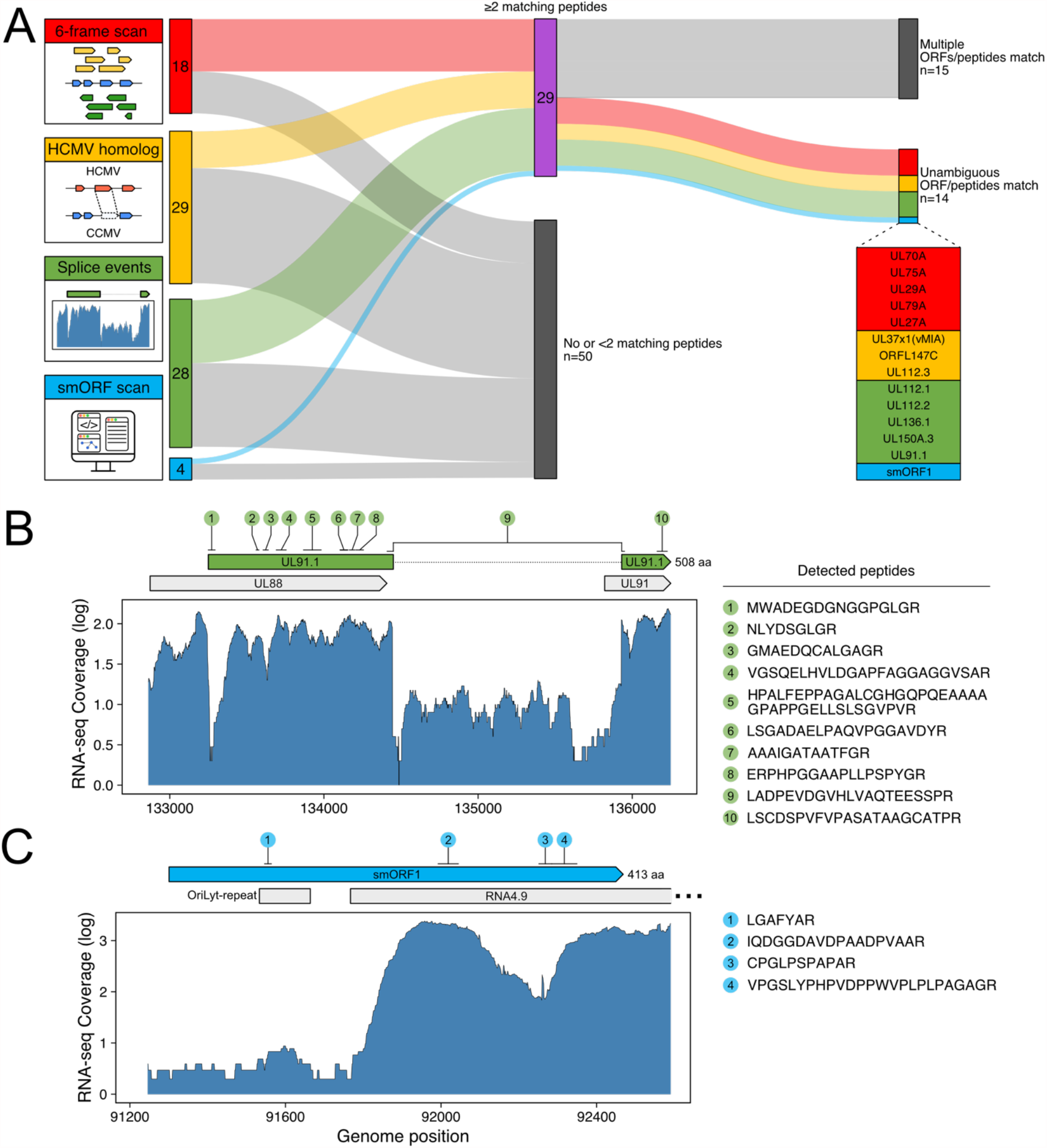
Re-evaluation of the CCMV coding potential. (A) To annotate potential novel coding sequences, the CCMV genome was scanned for unknown six-frame translation products, for homologs of non-canonical HCMV ORFs (6), for splice variants and for small ORFs (smORFs). To validate the candidate ORFs (numbers are indicated), proteomic data of CCMV-infected cells was analyzed for matching peptides. In total, 14 novel gene products were unambiguously identified by at least two unique peptides. (B) UL91.1 as an example of a newly found non-canonical CCMV ORF. RNA sequencing (RNA-seq) of infected cells detected a splicing event between the UL88 and UL91 locus. Multiple peptides matched the annotated ORF sequence, including the N-terminal peptide and a peptide spanning over the exon/exon junction, thus confirming translation of this splice variant. (C) smORF1 is another unique ORF with yet unknown function that overlaps the lytic replication origin of CCMV including the 5’-end of the long non-coding RNA4.9.

An example of a newly identified CCMV gene product is UL91.1 that originates from splicing of a 402 codons long alternative reading frame within the UL88 locus to codon 37 of the UL91 ORF, sharing the 107 C-terminal codons with UL91 (Fig. 5B). In contrast, the newly identified 413 codons long smORF1, which is encoded within the lytic replication origin (Fig. 5C), has no matching sequences in HCMV or other organisms, indicating this ORF to be unique to CCMV. Other newly annotated coding sequences (UL29A, UL70A, UL79A) are partly conserved in HCMV, but the corresponding ORFs are defective and interspersed with stop codons (Fig. S8), indicating that this group of genes was lost during HCMV evolution.

Besides the 14 newly annotated ORFs, we found peptides in our proteomic data set matching N-terminal sequences of UL48A, UL97, UL117 and US19 but spanning over the canonical methionine start codon, indicating N-terminal extensions of the respective gene products. Further, we annotated splicing sites, four long non-coding RNAs and nine micro-RNAs to CCMV-Phan9-BAC. An annotated genomic sequence file is available under the NCBI accession number MZ151943.

### Temporal profiling of CCMV protein expression

HCMV gene expression follows a strict temporal order with each protein falling into one of five distinct kinetic classes (Weekes et al., 2014). To determine whether this level of regulation is conserved in CCMV infection, we determined the kinetic profiles of each identified CCMV protein (n=142) and clustered these by the k-means method. The optimal number of clusters lies at five for our CCMV data set (Fig. S9). This was in congruence with HCMV (Weekes et al., 2014), indicating that the overall temporal organization of the viral gene expression cascade is conserved between both viruses. Following the nomenclature used for temporal profiles (TP) of HCMV protein expression (Weekes et al., 2014), we termed these clusters TP1-TP5 (Fig. 6). Similar to HCMV, TP1 and TP4 clusters of CCMV contained the lowest number of proteins and TP5 the highest.

**FIG 6.**
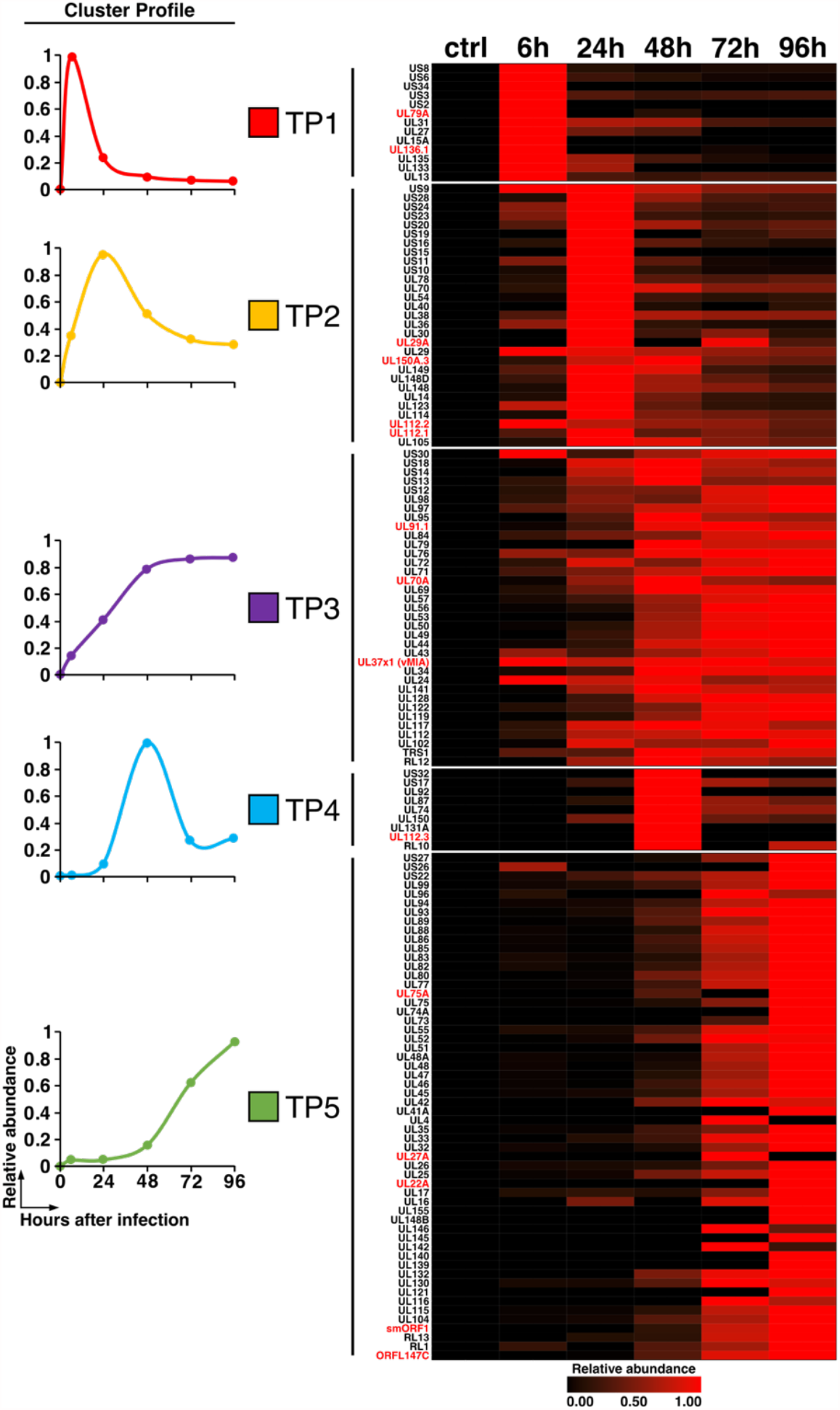
Temporal classes of CCMV protein expression. CCMV-infected cells were harvested at the indicated times post infection and subjected to a mass spectrometry-based proteomic analysis. K-means clustering classified viral gene expression into five temporal profiles (TP). The left-hand panel displays the average cluster profiles of TP classes 1-5. On the right hand, the expression profiles of individual CCMV proteins are plotted as a heatmap.

We reasoned that viral proteins that are functionally conserved between HCMV and CCMV should be produced with similar kinetics. Therefore, we compared the individual assignment of homologous viral gene pairs to the temporal clusters (Fig. S10). Genes assigned to the late expression profile (TP5) showed the largest overlap (65% of HCMV, 83% of CCMV proteins) and most non-overlapping genes of TP5 were assigned to the similarly shaped early-late profile TP3 (19% of HCMV, 10% of CCMV TP5 proteins). This indicates that molecular processes driving the accumulation of structural genes are largely conserved between both viruses. In contrast, only one gene of the TP4 clusters overlapped between HCMV and CCMV, suggesting that this small group of genes, with peak expression levels at 48 h post infection, may have less conserved functions.

### Regulation of host gene expression by CCMV

We next aimed to complete our gene expression analysis of CCMV-infected cells by evaluating the level of host gene regulation. To allow direct comparison to HCMV, we included in our analysis a previously published dataset of HCMV-infected cells (Nightingale et al., 2018). First, we calculated the changes in protein abundances induced by CCMV and HCMV and aligned the datasets based on shared Uniprot IDs (see table S5). Then, we determined the top 3% most up- and down-regulated host factors at each time point of CCMV and HCMV infection (Fig. 7A). To identify functionally related proteins enriched in the individual gene sets, we conducted a gene ontology (GO) analysis (Zhou et al., 2019). At late times of infection, both viruses showed a strikingly similar pattern in the down-regulation of pathways associated with actin cytoskeleton and extracellular matrix organization, cell adhesion, Rho GTPase, growth factor and cytokine signaling as well as programmed cell death (Fig. 7B, lower panel), suggesting that previously reported HCMV gene functions (Brune and Andoniou, 2017; Jarvis et al., 2006; Reinhardt et al., 2006; Stanton et al., 2014, 2007) are conserved in CCMV. At early times of infection, HCMV and, to a lesser extent, CCMV interfered with the expression of protein clusters associated with dNTP and amino acid metabolism as well as vesicle-mediated transport. Conversely, both viruses showed an early induction of proteins involved in mRNA processing and DNA repair (Fig. 7B, upper panel). Moreover, protein folding and a broad spectrum of metabolic processes were similarly enriched by HCMV and CCMV at later stages of infection. Of note, several GO terms describing mitochondrial processes, known to be important for HCMV infection (Combs et al., 2020), are up-regulated solely by HCMV.

**FIG 7.**
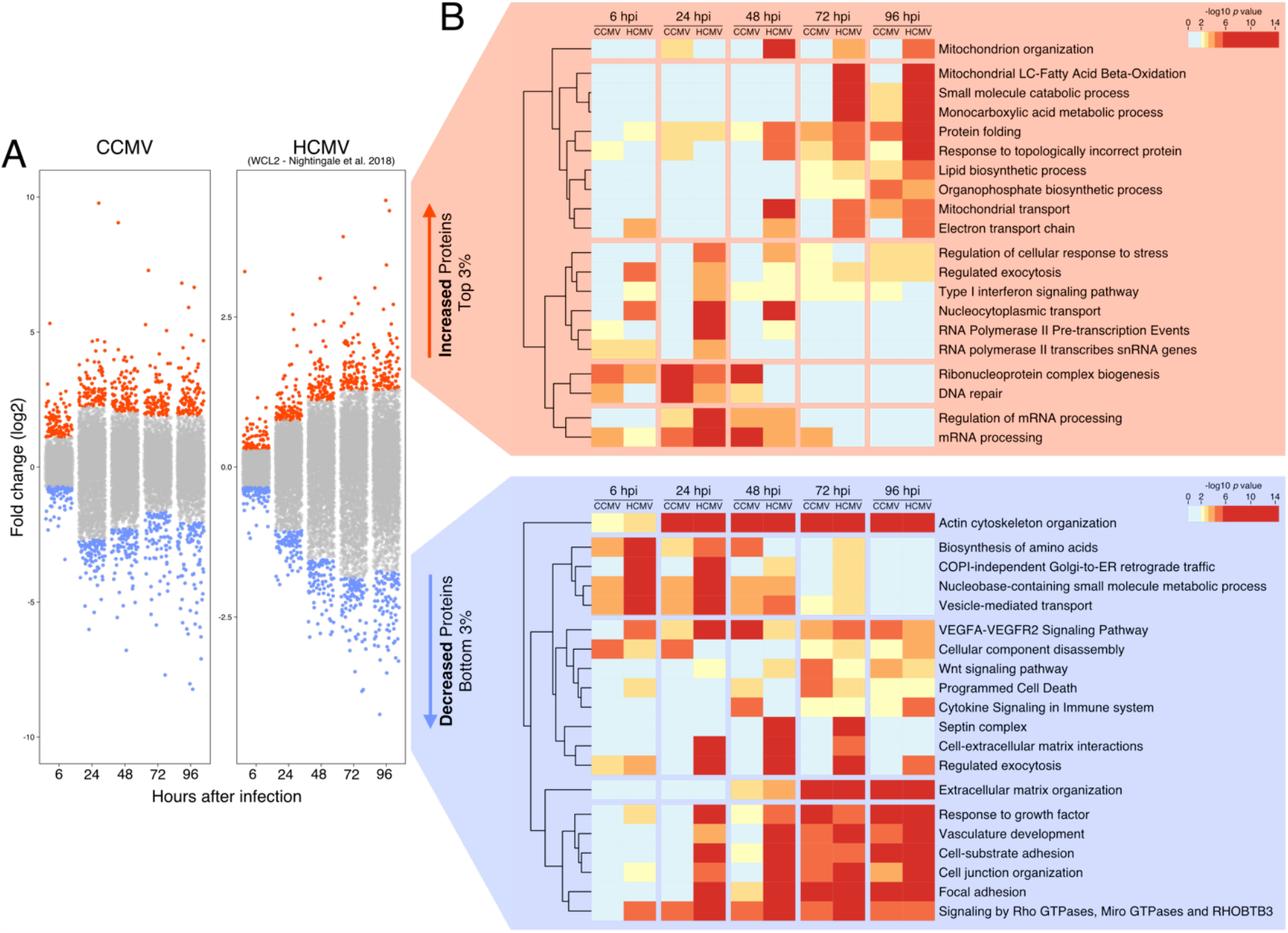
Gene ontology (GO) analysis of the most deregulated host proteins during CCMV and HCMV infection (HCMV data set WCL2 taken from Nightingale et al. 2018). (A) For each time point post infection, fold-changes in host protein expression were calculated relative to uninfected control cells (0 h). The top and bottom 3% of protein fold-changes were selected from each time point and used for multi-list GO enrichment analysis. (B) Heatmap showing the GO terms for strongly increased or depleted proteins during HCMV and CCMV infection (LC: long chain).

Next, we characterized the dynamics of mRNA and protein expression of 4,748 host genes during CCMV infection in order to distinguish between transcriptionally regulated host factors and those that are down-regulated at the protein level. We performed a k-means clustering analysis (Fig. S12) and found that the host gene expression profiles fall into seven distinct groups (Fig. 8). Exactly the same optimal number of clusters was found by Nightingale et al. when analyzing host mRNA and protein expression patterns in HCMV-infected fibroblasts (Nightingale et al., 2018). Three clusters in our analysis show relatively stable transcript levels but decreased protein amounts (Fig. 8, left panel) at either early (cluster 6), late times (cluster 5) or throughout infection (cluster 7). Such patterns are typical for virus-induced degradation of proteins hindering the infectious process. In fact, clusters 5 and 7 include the cyclin-dependent kinase inhibitor CDKN1A and the E3 ubiquitin ligase NEDD4 (Fig. 8, right panel), which are known targets of protein degradation by HCMV (Chen et al., 2001; Nightingale et al., 2018). Cluster 6 includes MYO18A which shows an initial degradation but a rapid increase at later stages of infection, which is in line with MYO18A being repurposed for virus assembly and egress during HCMV infection (Jean Beltran et al., 2016).

**FIG 8.**
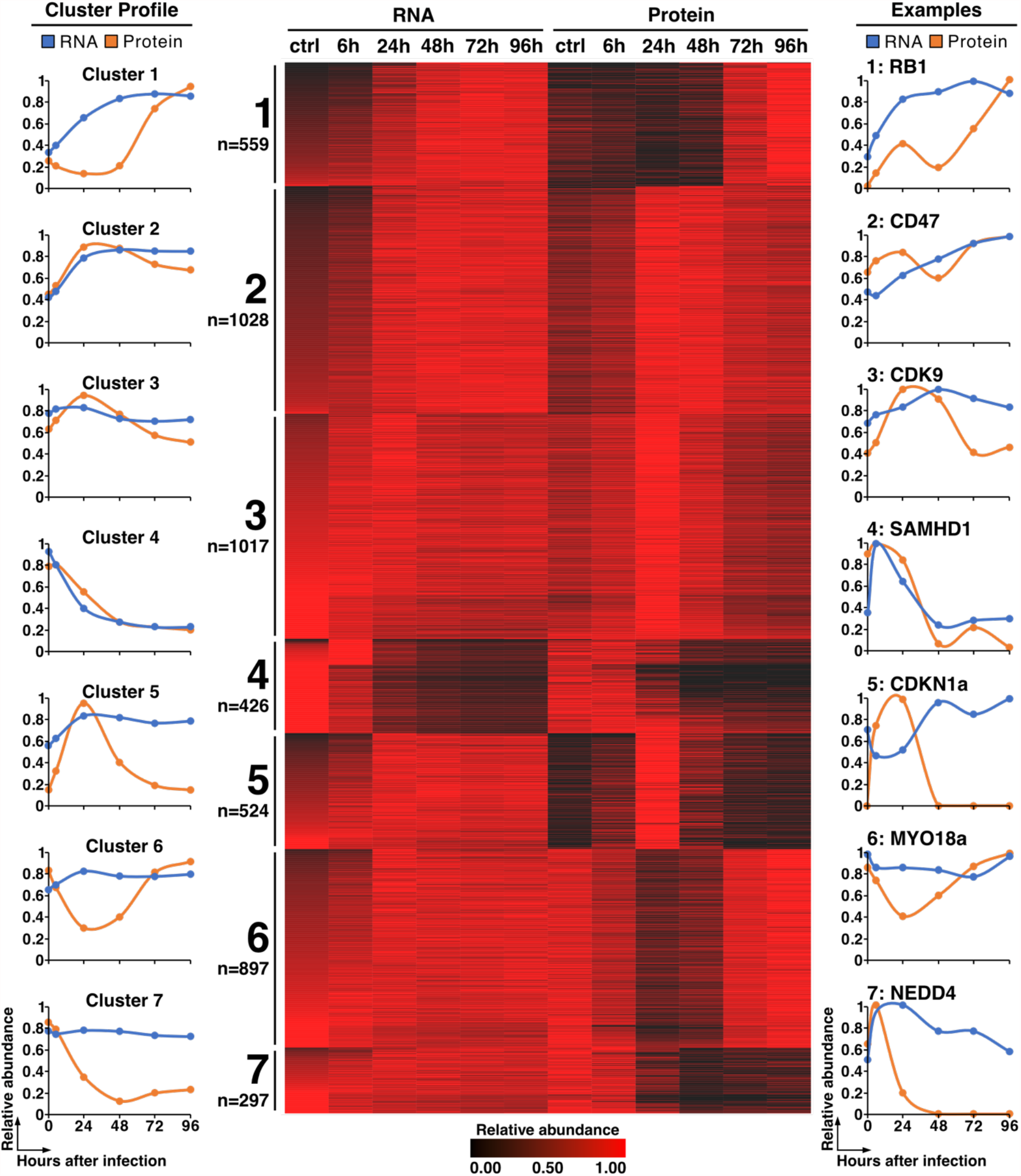
Comparative analysis of CCMV-induced changes in host gene expression at the transcript and protein level. At the indicated time points total RNA and protein were prepared from CCMV-infected cells and analyzed by RNA sequencing and tandem mass spectrometry respectively. K-means clustering of 4,748 transcripts and proteins identifies seven individual clusters representing different modes of host gene regulation during CCMV infection. Shown are the average mRNA and protein profiles of each cluster (left panel), a heatmap of the corresponding gene expression data (middle panel) and selected representative profiles from each cluster (right panel).

Two other clusters contain genes whose protein levels closely parallel mRNA expression, indicating transcriptional activation (cluster 2) and repression (cluster 4), respectively. 49% of transcriptionally induced and 56% of transcriptionally silenced genes in CCMV infection are also present in the corresponding clusters from HCMV-infected cells (Fig. S13). This suggests that the regulation of host gene transcription is generally well conserved between both viruses. An example of a transcriptionally repressed gene during CCMV infection is the dNTP hydrolase SAMHD1 which acts as a restriction factor for many viruses including HCMV (Businger et al., 2019; Kim et al., 2019). Thus, our database of mRNA and protein profiles in CCMV infection can function as a valuable resource for identifying host targeting mechanisms shared with HCMV and sets the framework for a comparative assessment of conserved virus-host interactions.

## DISCUSSION

Animal CMVs and their BAC clones have been utilized for decades for *in vivo* and *in vitro* models and thereby contributed immensely to the understanding of pathogenesis, host immune response and underlying molecular mechanisms of CMV infections. As the closest relative to HCMV, the chimpanzee CMV provides a link between the human and other animal CMV species and holds a unique potential to serve as a model system for HCMV infection. Here, we established a BAC tool based on the genome of the CCMV Heberling strain, making CCMV for the first time readily accessible to reverse genetic approaches. The system-level characterisation of RNA and protein expression led to the identification of novel viral gene products and highlighted the many shared attributes of CCMV and HCMV gene regulation, providing resource and guidance for the usage of CCMV as a model.

The straightforward approach to integrate the BAC cassette into the viral genome by homologous recombination proved to be a reliable cloning strategy for CCMV as it has been for HCMV genomes before (Borst et al., 1999; Christian Sinzger et al., 2008; Tischer and Kaufer, 2012). However, recombinant viral genomes exceeding the natural genome length provoke deletions or genomic rearrangements to accommodate the viral genome into the capsid structure. Indeed, deletions of in total 8.8 kbp adjacent to the BAC cassette were detected in all our initially selected BAC clones (Fig. 1) and similar deletion events occurred during the generation of TB40-BAC4 (Christian Sinzger et al., 2008). While this sets a limitation on BAC-capturing the full length viral genome at first instance, deleted sequences can be seamlessly reinstated later by BAC mutagenesis protocols as it has been done in this study and for TB40-BAC4 (Sampaio et al., 2017; Christian Sinzger et al., 2008). In the future, approaches based on the assembly of synthetic or separately cloned subgenomic fragments may reduce the risk of such unwanted deletion and recombination events and become attractive alternatives to traditional BAC cloning procedures (Oldfield et al., 2017; Thi Nhu Thao et al., 2020).

Whole genome sequencing of the Heberling strain revealed two polymorphic loci, US26-28 and UL48-55, within the CCMV genome and a large set of the variations carried over in different combinations to the five sequenced BAC clones (Fig. 2A, Fig. S6). There are several possible explanations for the origin of this genetic diversity. As CCMV Heberling was never plaque-purified after isolation from the donor animal, some of the observed variations may reflect the pre-existing viral intra-host diversity. For instance, the UL55 gene of HCMV, encoding glycoprotein B (gB), is also polymorphic and occurs in mixed genotypes in infected individuals (Renzette et al., 2011). The positions of non-synonymous mutations found in CCMV-UL55 directly correspond to the most variable gB regions in HCMV (Fig. S7A), including antigenic domains 1, 2 and near the furin cleavage site (Foglierini et al., 2019; Pignatelli et al., 2004). This suggests that the different UL55 genotypes of CCMV represent neutralizing antibody escape variants originating from the original chimpanzee sample.

Another source of genetic diversity are evolutionary processes like selection and genetic drift, acting on CCMV during *in vitro* culture. The CCMV isolate not only had to adapt to a human host cell environment but was selected for at least twenty passages for cell-free growth on fibroblasts (Davison et al., 2003), without any selective pressure by the immune system. Well known adaptations of HCMV to the cultivation on fibroblasts are disruptive mutations in the UL128 locus, the RL13 gene and the ULb’ region (Wilkinson et al., 2015). While CCMV Heberling contains a frameshift mutation in UL128, facilitating the growth on fibroblasts but rendering the virus unable to infect endothelial and epithelial cells (Fig. 3), RL13 and ULb’ genes lack any obvious frameshift or translation terminating mutations. Adaptation processes of primate CMVs to human cells can occur in a rapid fashion facilitated by recombination events (Child et al., 2021) or by positive selection of advantageous genetic variants (Mozzi et al., 2020). UL48, which exhibits the second highest number of non-synonymous polymorphisms among all CCMV genes (Fig. 2A, Table S1), has been predicted to be one of the factors driving the evolutionary adaptation to the human host (Mozzi et al., 2020). The fact that one half of all non-synonymous polymorphisms in CCMV-UL48 cluster to a narrow central region whereas none is located in the N-terminal deubiquitinase domain (Fig. S7B), argues that non-enzymatic functions of this inner tegument protein might be involved in such adaptation processes (Kim et al., 2016; Yu et al., 2011). In contrast to the UL48-UL55 region, which encompasses essential core genes, the overall high frequency of nucleotide variants across the US region (Fig. 2A, Fig. S6) is likely due to the fact that US genes are dispensable for virus growth *in vitro* (Dunn et al., 2003; Hein and Weissman, 2021; Yu et al., 2003) and therefore tolerate mutations.

By re-evaluating the coding potential of CCMV we identified 14 novel ORFs that could be confirmed by mass spectrometry (Fig. 5A, Table S2). Among the newly annotated genes, several are unique to CCMV, including the UL91 splice variant UL91.1 and the oriLyt-spanning smORF1 (Fig. 5B-C). UL91 is an integral part of the viral pre-initiation complex (vPIC) that governs the transcriptional activation of “true” late genes (Gruffat et al., 2016). The UL91 N-terminus (residues 1-70) is conserved among primate CMVs and its integrity is essential for vPIC function (Omoto and Mocarski, 2013). Considering that residues 1-36 of UL91 are lacking in UL91.1, it is highly unlikely that the UL91 activity is preserved. Possibly, the shared C-terminus enables UL91.1 to act as a regulator of vPIC, either by competing with UL91 binding or by recruitment of the large, intrinsically disordered N-terminal domain encoded by UL91.1 exon 1. Although UL91.1 splice sites are conserved in RhCMV and HCMV (Gatherer et al., 2011), a UL91.1 exon 1-related ORF is completely lacking in these viruses. This stands in contrast to UL29A, UL70A and UL79A which, although being unique to CCMV, have homologous, yet fragmented and non-functional ORFs at related positions of the HCMV genome (Fig. S8). Hence, it appears that UL91.1 has been newly acquired by CCMV while UL29A, UL70A and UL79A derive from common ancestors that were lost during HCMV evolution.

The finding that smORF1 overlaps with the oriLyt of CCMV was surprising, as lytic replication origins of herpesviruses are typically devoid of protein coding sequences and instead regulated by non-coding transcription (Boldogkői et al., 2019). The long-noncoding RNA4.9 of HCMV activates viral DNA replication by R-loop formation, which exposes the oriLyt to the viral single-stranded DNA-binding protein UL57 (Tai-Schmiedel et al., 2020). The abundant expression of an RNA4.9 homolog from the oriLyt region of CCMV (Fig. 5C) indicates that this mechanism is conserved, despite the presence of smORF1. Concerning the potential conflicts between transcription and replication processes (Gómez-González and Aguilera, 2019), the kinetics of smORF1 expression (TP5, Fig. 6, Table S4) may indicate that smORF1 transcription impacts the initiation of viral DNA synthesis at late stages of CCMV infection.

The temporal analysis of CCMV protein expression reveals that the majority of gene products fall in the same kinetic classes as their HCMV orthologues (Fig. S10A) and most of the observed class switches can be considered as conservative (Fig. S10B). Nevertheless, a few orthologs show drastic differences in their kinetic profiles (Fig. S11). This latter group includes UL148 which has early expression kinetics in CCMV (TP2) but late kinetics (TP5) in HCMV infection. HCMV-UL148 has two distinct functions: it promotes the intracellular retention of CD58, thereby impeding the detection of infected cells by the immune system (Wang et al., 2018); and it causes large-scale remodeling of the endoplasmic reticulum (ER), thereby protecting the viral glycoprotein O (UL74) from ER-associated degradation (Nguyen et al., 2020, 2018). Since only the immune evasion but not the ER remodeling function is conserved in CCMV-UL148 (Nguyen et al., 2020), it is conceivable that its earlier expression reflects the lack of ER remodeling function. In fact, HCMV-UL74 expression (TP5) parallels that of UL148 (Fig. S10-11), suggesting that the temporal shift of HCMV-UL148 expression has evolved to ensure proper maturation of viral envelope proteins during the assembly phase of infection. This would support a general scenario, where distinct expression kinetics indicate functional divergence of viral orthologs.

Our systems-level overview of host dynamics during CCMV infection and comparison to HCMV data from previous studies highlights many common themes of CMV infection (Fig. 7B) and thus further underscores the modelling potential of CCMV. For example, clusters 5 and 7 contain host factors with steady mRNA levels but a rapid decrease in protein abundances as infection progresses (Fig. 8), suggesting these factors are targets of virus-induced protein degradation. Clusters 5 and 7 include NEDD4, NEDD4L and ITCH, members of the NEDD4 family of E3 ubiquitin ligases that recognize a PPXY docking motif on their protein substrates (Ingham et al., 2004). HCMV-UL42 employs two PPXY motifs in its N-terminus to interact with NEDD4 family members and causes their proteolysis (Koshizuka et al., 2018, 2016; Nobre et al., 2019), possibly to protect other viral proteins from NEDD4-mediated ubiquitination (Soh et al., 2020). The presence of three PPXY motifs in the CCMV-UL42 N-terminus supports the notion that NEDD4 degradation is a shared strategy of herpesviruses (Koshizuka et al., 2018).

Our searchable database with transcript and protein dynamics of 4,748 cellular genes allows a quick assessment of further relevant virus-regulated host factors and signaling pathways (Table S6). This can provide insights into host subversion strategies that are shared between CCMV and HCMV. Such comparative analyses are instrumental for the identification of critical nodes of virus-host interaction. In addition, the CCMV system offers a versatile genetic tool to dissect essential functions and infection mechanisms of HCMV by gene or domain swap experiments. All these comparative approaches are greatly facilitated by the fact that both viruses can be analyzed in the same cellular background. Thus, although CCMV research is limited to in vitro systems (Knight, 2008), the close phylogenetic relationship of CCMV to HCMV offers some key advantages over other CMV species that, with the help of our in-depth characterized virus clone, can be exploited in future studies.

## MATERIALS AND METHODS

### Cell culture and viruses

Human embryonic lung fibroblasts (Fi301 – obtained from the Institute of Virology, Charité Berlin, Germany) were maintained as previously described (Zydek et al., 2010). HUVEC cells were kindly provided by Andrea Weller (Center for Cardiovascular Research (CCR), Charité Berlin, Germany) and maintained as previously described (Stangl et al., 2001). ARPE-19 cells (ATCC^®^ CRL-2302) were cultured in DMEM 10% FCS with 20 mM L-Glutamine and 45 ug/ml gentamicin. The Chimpanzee Cytomegalovirus Heberling strain was kindly provided by Gary S. Hayward (Johns Hopkins School of Medicine, Baltimore, MD, USA). For preparation of viral stocks, CCMV Heberling strain was added to Fi301 cells at an MOI of 0.1 and infected cells were kept in subculture until all cells showed a cytopathic effect. The medium was then changed to DMEM supplemented with 2% FCS, 20 mM L-Glutamine and 45 ug/ml Gentamicin (harvest medium). Viral supernatant was harvested 3-4 days post medium change, centrifuged at 3000 rcf for 15 minutes and stored as 2 ml single-use aliquots at -80°C. For infection experiments that required a high MOI, virus was concentrated prior infection by ultracentrifugation at 30.000 rcf for 40 minutes at 25°C on a 20% sucrose cushion. Virus pellet was then resuspended in harvest medium and added to the cells at the desired MOI.

### BAC cloning

The BAC donor plasmid pEB1097 (kindly provided by Eva Borst, Hannover) contains a BAC cassette flanked by homology arms suited to the US1-2 and US6-7 sequences of HCMV strain AD169 (Borst et al., 1999). The homology arms of pEB1097 were replaced with the corresponding genomic regions of CCMV Heberling (coordinates left homology arm: 204.201-205.187 bp, right homology arm: 208.104-209.072 bp) via NEBuilder HiFi DNA Assembly Cloning (New England Biolabs). The resulting vector was named pEB1097_CCMV (Fig. S1). Recombinant virus was generated by spontaneous recombination of the BAC cassette and the CCMV genome in infected cells. First, the BAC cassette including the homology arms was PCR-amplified from pEB1097_CCMV, using the primer pair 69_CCMV_donor_PCR_fw and 70_CCMV_donor_PCR_rev. 1 µg of the amplicon was transfected into 10^6^ Fi301 cells by Amaxa nucleofection, using the “Primary Fibroblast” kit and pulse program A-023 (Lonza). The day after, cells were infected with CCMV Heberling at an MOI of 1. 4 days post infection, virus containing supernatant was transferred to a fresh culture of Fi301 cells. From then on, selection was applied by adding 200 µM xanthine and 200 µM mycophenolic acid to the culture medium. After three rounds of virus passages with selection, the circular virus DNA was extracted from infected cells using the Hirt method (Hirt, 1967) and electroporated into E.coli ElectroMAX DH10B Cells (Invitrogen). Transformants were selected on LB agar plates with 12.5 µg/ml chloramphenicol and colonies were picked for further characterization.

### BAC mutagenesis

CCMV BAC mutants were created by en passant mutagenesis as previously described (Tischer et al., 2010). Authentic recombination events were verified via PCR analysis and Sanger sequencing. Overviews of the BAC cloning strategies are shown in supplementary figures S2-S4 and supplementary table S3 lists all used oligonucleotides.

### Reconstitution of CCMV BAC virus

BAC-DNA was prepared using the NucleoBond Xtra-Midi kit (Macherey-Nagel) according to the manufacturer’s instructions. A mixture of 2 µg of BAC-DNA, 2 µg pcDNA-pp71-flag and 1 µg of pBRep-Cre (Wolfram Brune, Hamburg, Germany) was electroporated into Fi301 cells using the Amaxa nucleofector program A-23 (Lonza). Plaque formation was checked over the course of 14 days, cells were kept in subculture and once all cells showed a cytopathic effect, medium was changed to harvest medium and viral supernatant was harvested 3 days post medium change.

### Viral DNA extraction

For isolation of viral DNA, viral particles were ultra-centrifuged at 30,000 rcf for 30 minutes at 25°C. Supernatant was discharged and the pellet was resuspended in 100 µl Tris buffer containing proteinase K and incubated overnight at 56°C. DNA was then extracted via Phenol-Chloroform extraction using Roti-Phenol (Carl Roth).

### Flow cytometry

CCMV infectious titers were assessed based on the ability to induce IE1 expression 24 hours after infection. Infected cells were harvested by trypsinization, fixed and permeabilized by ice-cold absolute ethanol and incubated for at least 10 minutes at 4°C. Cells were then stained with Alexa Fluor 488-conjugated anti-IE1/IE2 (MAB810X) antibody overnight at 4°C. Flow cytometry was done on a FACSCanto II flow cytometer (BD Biosciences) using FACSDiva (BD Biosciences) and FlowJo (FlowJo LLC) software packages. Cellular debris, cell doublets and aggregates were gated out of analysis.

### CCMV DNA sequencing and variant analysis

DNA from CCMV BAC clones and the CCMV Heberling stock was sequenced at Eurofins Genomics, Ebersberg with Illumina 2×150 bp paired end-reads. Eurofins variant analysis included read-mapping against the given reference genome, detection and annotation of single nucleotide variations (SNVs) and InDels as well as the allocation of their effects on protein level. CCMV genome region, ORFs, DNA sequencing coverage and positions of SNVs were plotted using Circos 0.69-9.

### Total RNA sequencing

Confluent Fi301 cells were infected in triplicates with Phan9-BAC-derived CCMV using an MOI of 5. Samples were harvested by resuspension in Trizol (Thermo Fisher Scientific). RNA extraction, library preparation, total RNA sequencing and data processing were performed as previously described (Gonzalez-Perez et al., 2021). Splice junctions were identified using Geneious Prime software.

### Mass spectrometry

Infection conditions were the same as for the sample preparation for RNAseq. Infected cells were harvested by removing the medium and scraping off the cell layer in 1 ml PBS. For lysis, the cell pellet was resuspended in 200 µl of lysis buffer: 7 M Urea, 50 mM triethylammonium bicarbonate (pH 8.5), 1% Triton-X-100, 5 mM Tris(2-carboxyethyl)phosphine, 30 mM chloroacetamide, complete mini EDTA free protease inhibitors (Roche), 2 µl Benzonase (Merck-Millipore) and sonicated for 45 min (intervals: 20 s on, 40 s off) using a Bioruptor (Diagenode). After clearing the lysates, protein was isolated using methanol/ chloroform precipitation, according to standard protocols (Wessel and Flügge, 1984). The precipitates were resuspended in digestion buffer, consisting of 50 mM triethylammonium bicarbonate (pH 8.5), 1% sodium deoxycholate, 5 mM Tris(2-carboxyethyl)phosphine, 30 mM chloroacetamide, Trypsin (1:25, w/w), Lysyl Endopeptidase (1:100, w/w), and digested overnight at room temperature. Peptides were desalted using two disks of C18 material embedded in Stage Tips (Rappsilber et al., 2003) and stored at 4°C until LC-MS/MS measurement.

The LC/MS analysis was performed using a Thermo Scientific Dionex UltiMate 3000 system connected to a µPAC^™^ Trapping column (PharmaFluidics). The mobile phase A contained 2% acetonitrile and 0.05% Trifluoroacetic acid (TFA) in water and the mobile phase B contained 0.05% TFA in acetonitrile. For peptide separation a 200 cm µPAC^™^ micro pillar array column at 180 min gradient length was employed, with mobile phase A containing 0.1% formic acid in water, and mobile phase B 0.1% formic acid in acetonitrile. The flow rate was set to 750 nL/min from 4 % B to 20% B and 350 nL/min from 20 % B to 80 % B.

The MS1 scans were performed in the orbitrap using 120,000 resolution. Peptides were fragmented using higher-energy collision induced dissociation (HCD) with 30 % HCD collision energy. The MS2 scans were acquired in the ion trap with standard AGC target settings, an intensity threshold of 5e3 and maximum injection time of 40 ms. A 1 s cycle time was set between master scans.

Raw-files were searched using MaxQuant version 1.6.2.6 (Cox and Mann, 2008). Search parameters included two missed cleavage sites, fixed cysteine carbamidomethyl modification, and the variable modifications methionine oxidation, N-terminal protein acetylation as well as asparagine–glutamine deamidation. The “match between runs”, “iBAQ” (intensity-based absolute quantification), “second peptide” and “LFQ (label-free quantification)” options were enabled (Cox et al., 2014). Database search was performed using Andromeda, the integrated MaxQuant search engine, against a Uniprot database of homo sapiens proteins (downloaded 2020), the predicted amino acid sequences of the CCMV BAC and the proteins from a six frame translation of the CCMV genome (minimum sequence length: 15 amino acids) with common contaminants. False discovery rate was estimated based on target-decoy competition using a reverted database and was set to 1% at peptide spectrum match, protein and modification site level.

### Data analysis

For the multi-set gene ontology-enrichment analysis of the top 3% of proteins (highest and lowest log2 fold-change), the Metascape tool was applied (Zhou et al., 2019). Protein level data was matched to RNA level data based on the HGNC official gene symbol and were clustered using the k-means function in R (Gu, 2013). We determined the optimal number of clusters by calculating the summed distance of each protein from its cluster centroid and plotted these values against the corresponding numbers of clusters. The inflection point of this curve indicates the optimal number of clusters (“elbow method”). Plots were created using ggplot2 (Wickham, 2009) and pheatmap v.1.0.12 (https://CRAN. R-project. org/package= pheatmap).

## DATA AVAILABILITY

The mass spectrometry proteomics data have been deposited in the ProteomeXchange Consortium with the data set identifier PXD027434. The RNA sequencing data have been deposited in the NCBI GEO database, with the accession number GSE171149. An annotated genome sequence of CCMV-Phan9-BAC is available at GenBank under the accession number MZ151943.

## ACKNOWLEDGEMENTS

We are grateful to Benedikt Kaufer (Free University Berlin) for his helpful advice regarding the BAC cloning procedure. We thank Andrea Weller (Charité) and Eva Borst (Medizinische Hochschule Hannover) for providing valuable reagents. We also thank Iris Gruska and Barbara Vetter for their excellent technical assistance. This work was supported by a grant (#900005) from the Joachim-Herz-Stiftung to L.W. Q.V.P. was funded by a scholarship for postgraduate thesis projects (Charité Promotionsstipendium I) from the Charité Academics Grants Committee. BB acknowledges funding from DFG grant BO 5917/1-1.

