## Supplementary Figures S1-13 for "ENGINEERING, DECODING AND SYSTEMS-LEVEL CHARACTERIZATION OF CHIMPANZEE CYTOMEGALOVIRUS"

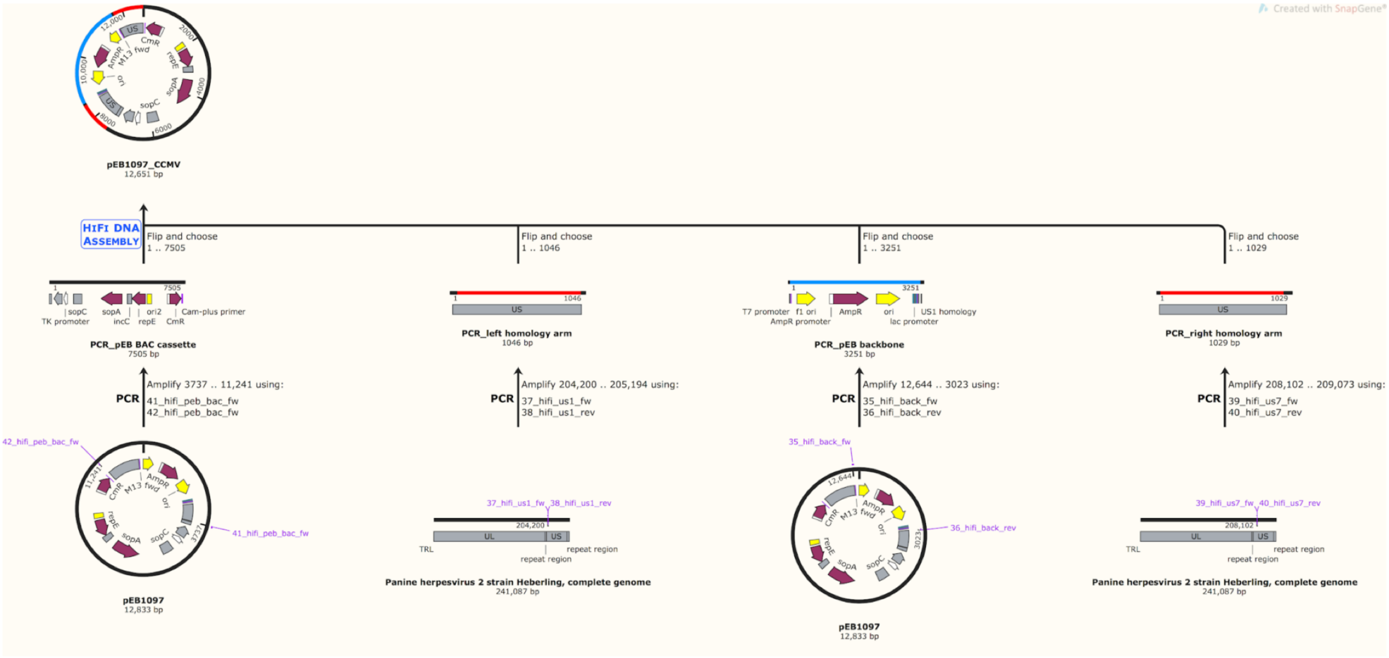


**FIG S1** Cloning scheme of the BAC donor vector pEB1097_CCMV. Homology arms for targeted integration into CCMV genome region US2-US6, BAC cassette and plasmid backbone sequences were PCR amplified and assembled via Gibson assembly (HIFI DNA assembly kit from NEB).

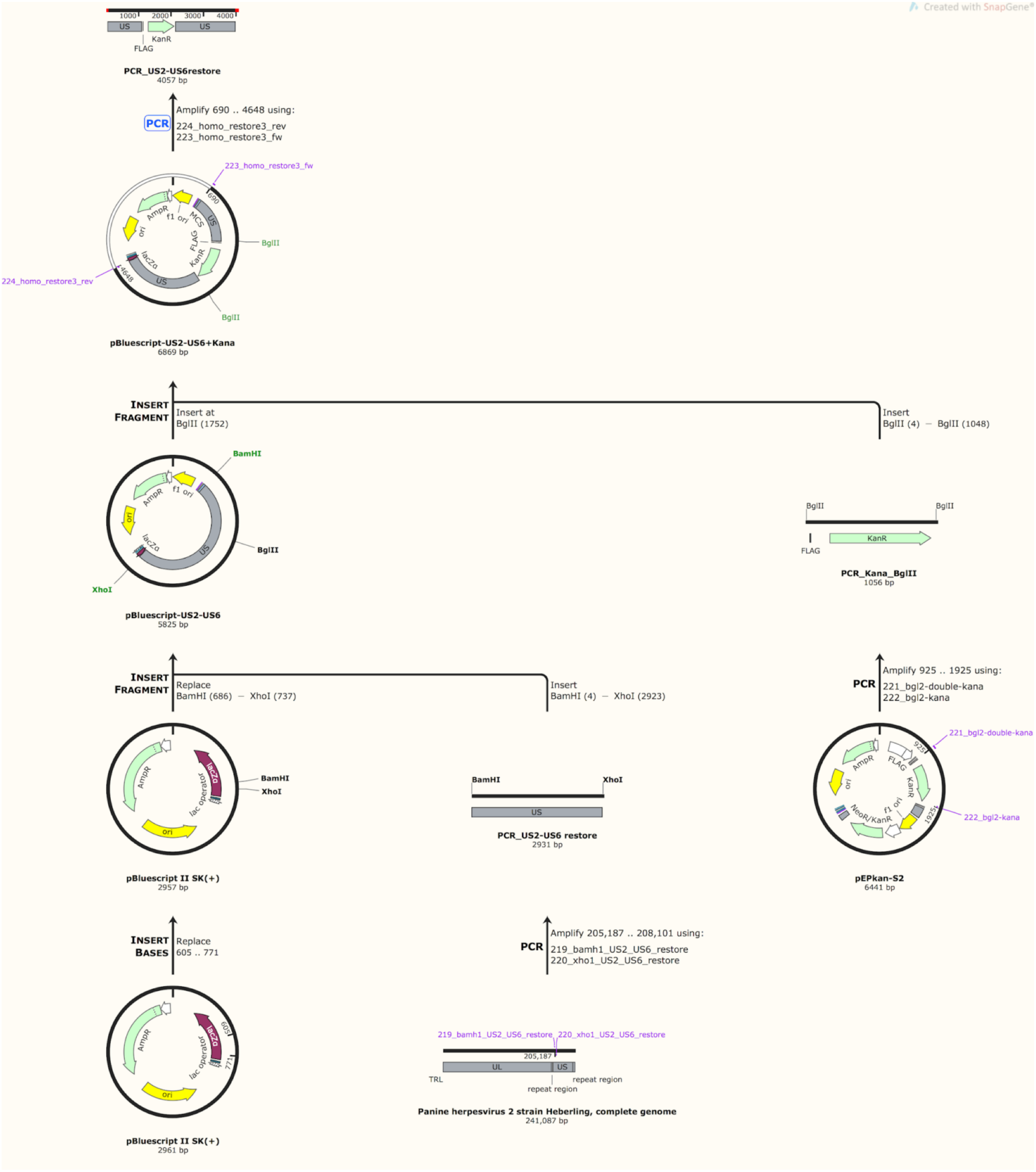


**FIG S2** Cloning of the transfer vector for the restoration of the deleted sequences on the left hand side of the BAC cassette in CCMV BAC 177.

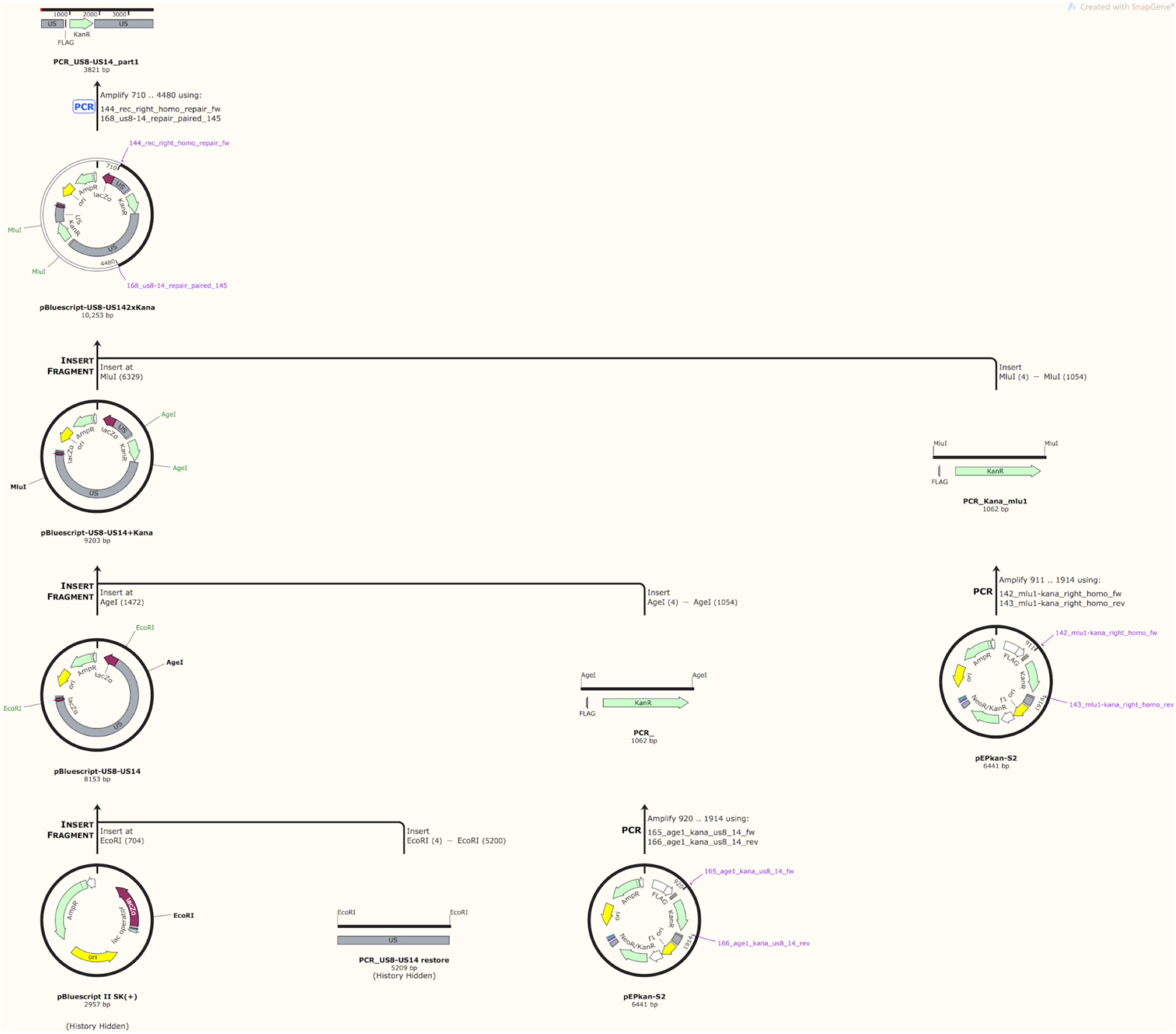


**FIG S3** Cloning of the transfer vector for the restoration of deleted sequences on the right hand side of the BAC cassette in CCMV BAC 177. Restoration was done in two parts: (A) Region US8-US10 (B) Region US11-US13

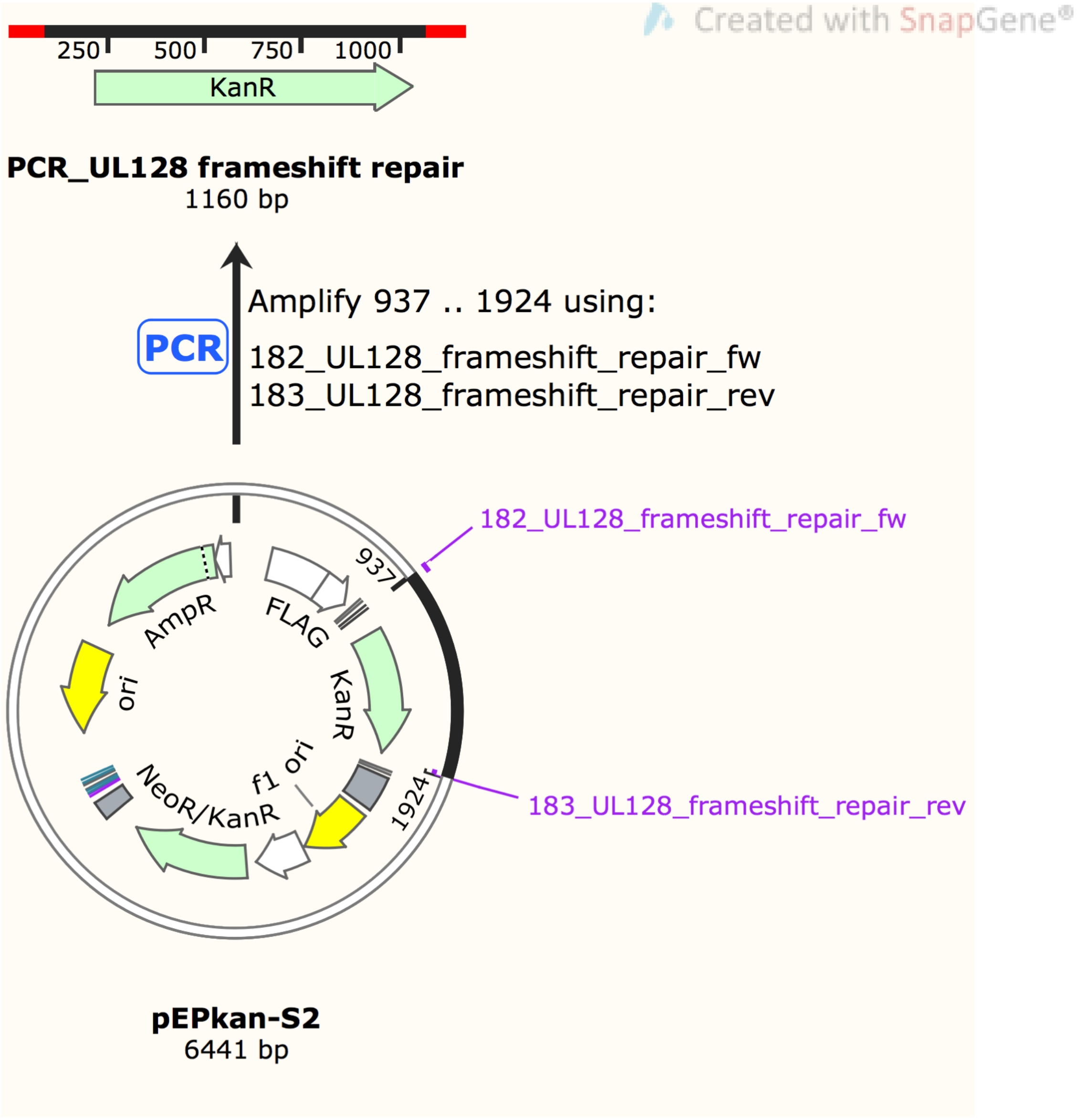


**FIG S4** PCR amplification scheme of the *en passant* recombination template for the UL128 frameshift repair.

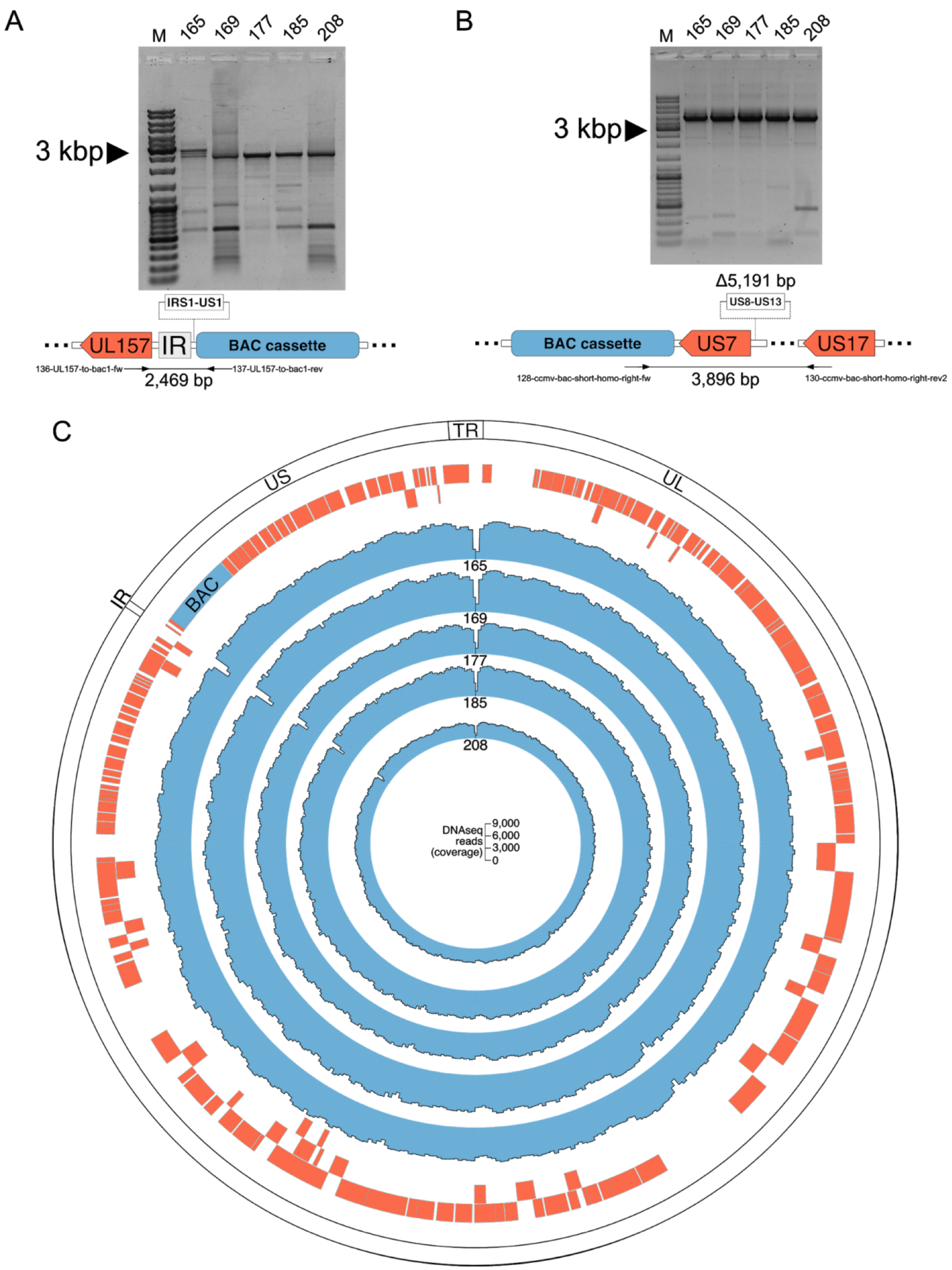


**FIG S5** Deletion events in the selected CCMV BAC clones. (A) PCR spanning over the left hand side of the BAC cassette integration site reveals deletion of IRS1 to US1 (3,639 bp). (B) PCR at the right hand side of BAC integration site reveals deletion of US8 to US13 (5,191 bp) in all clones. (C) Coverage plots show alignment of sequencing reads to adjusted reference genome. Alignment confirms deletion events and also reveals the same deletions in clone 165,169 and 208 at the left hand side of the BAC cassette. Tracks in inner circle shows the average coverage in 500 bp increments (blue) of each clone, CCMV ORFs (red) and genomic regions (white) are plotted in the outer tracks of the Circos plot.

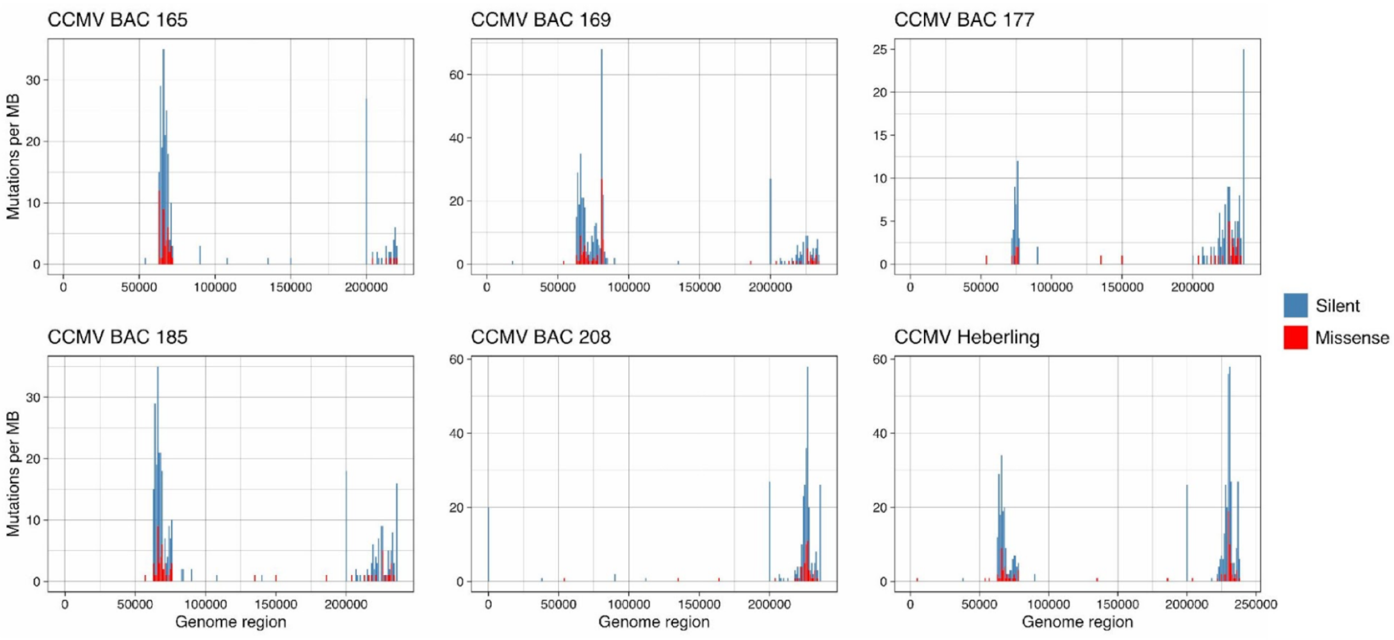


**FIG S6** Distribution of silent and missense mutations in selected CCMV BAC clones and the parental Heberling strain. Silent mutations are clustered at the two polymorphic loci with a higher prevalence than missense mutations.

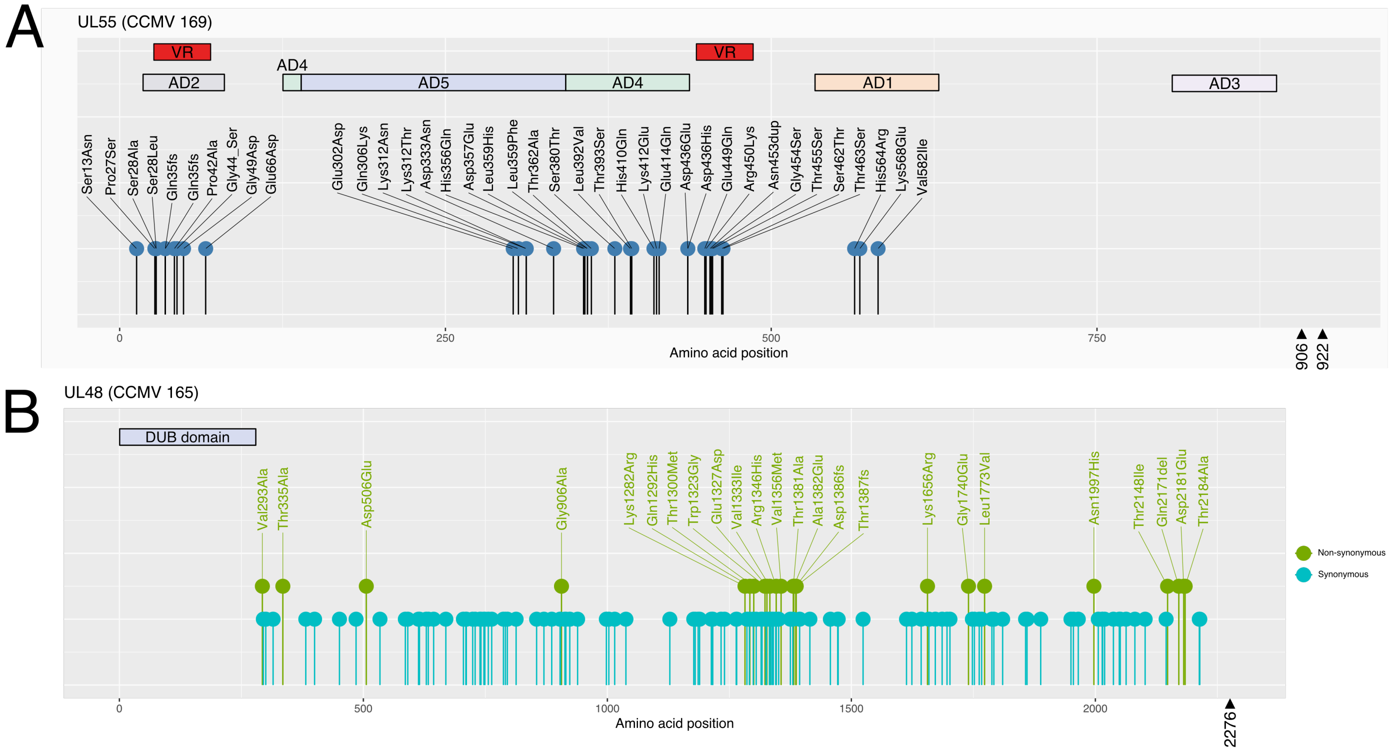


**FIG S7** Nature and position of amino acid (aa) changes in UL48 and UL55 (gB). (A) Mutation clusters found in UL55 (gB – 922 aa) of CCMV 169 with corresponding sequence areas of antigenic domains (AD) and variable regions (VR) described in HCMV-gB (902 aa). Positions of HCMV-gB domains taken from Foglierini et al. (2019) - *HCMV Envelope Glycoprotein Diversity Demystified.* (B) Amino acid changes in UL48 (2276 aa) of CCMV BAC 165. The corresponding sequence area of the HCMV-UL48 DUB domain is indicated.

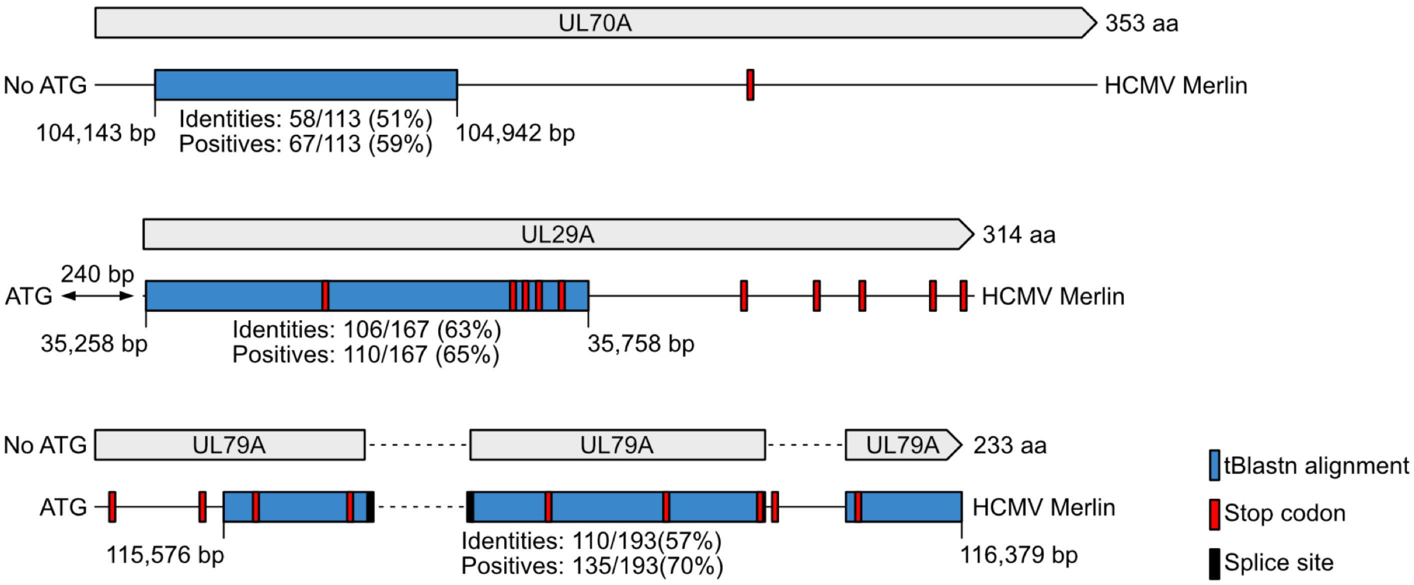


**FIG S8** Sequence regions in HCMV Merlin corresponding to the newly found CCMV ORFs. tBlastn scan identified sequence regions in HCMV Merlin (blue marked boxes) homologous to UL29A, UL70A and UL79A of CCMV. Genome coordinates, conserved splice sites, stop codons arising from frame-shift mutations and the percentage of amino acid identities and positives (amino acids that are either identical between the query and the subject sequence or have similar chemical properties) are indicated for the aligned sequence regions.

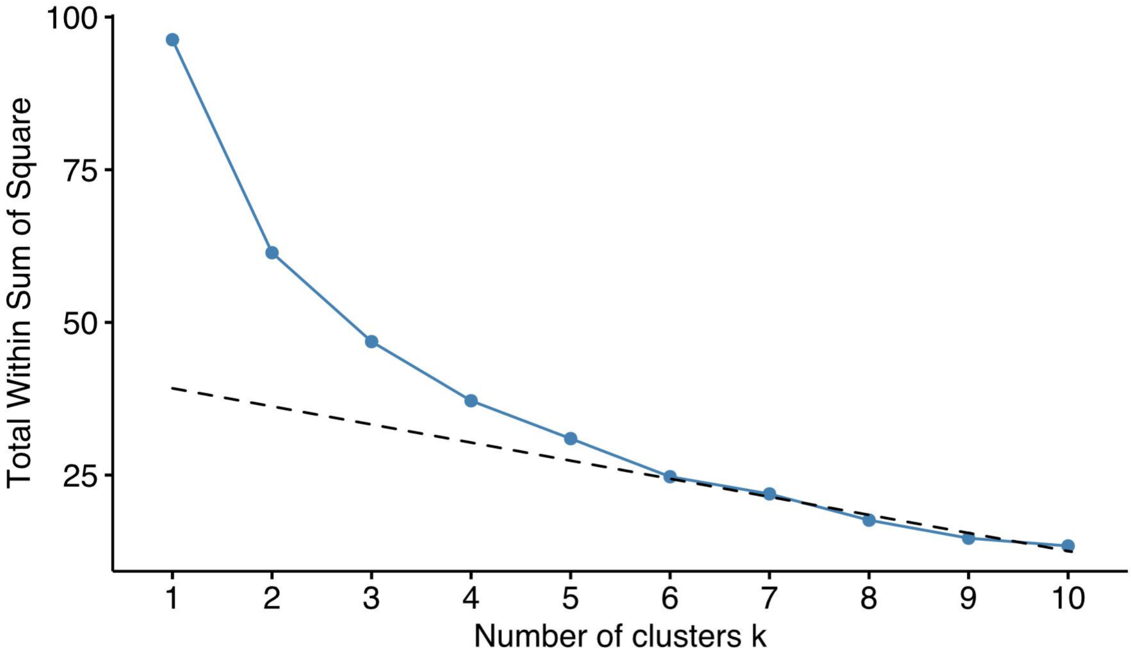


**FIG S9** Number of temporal classes of CCMV gene expression. The summed distance of each protein from its cluster centroid was calculated for one to 10 clusters and plotted. The point of inflection fell at five.

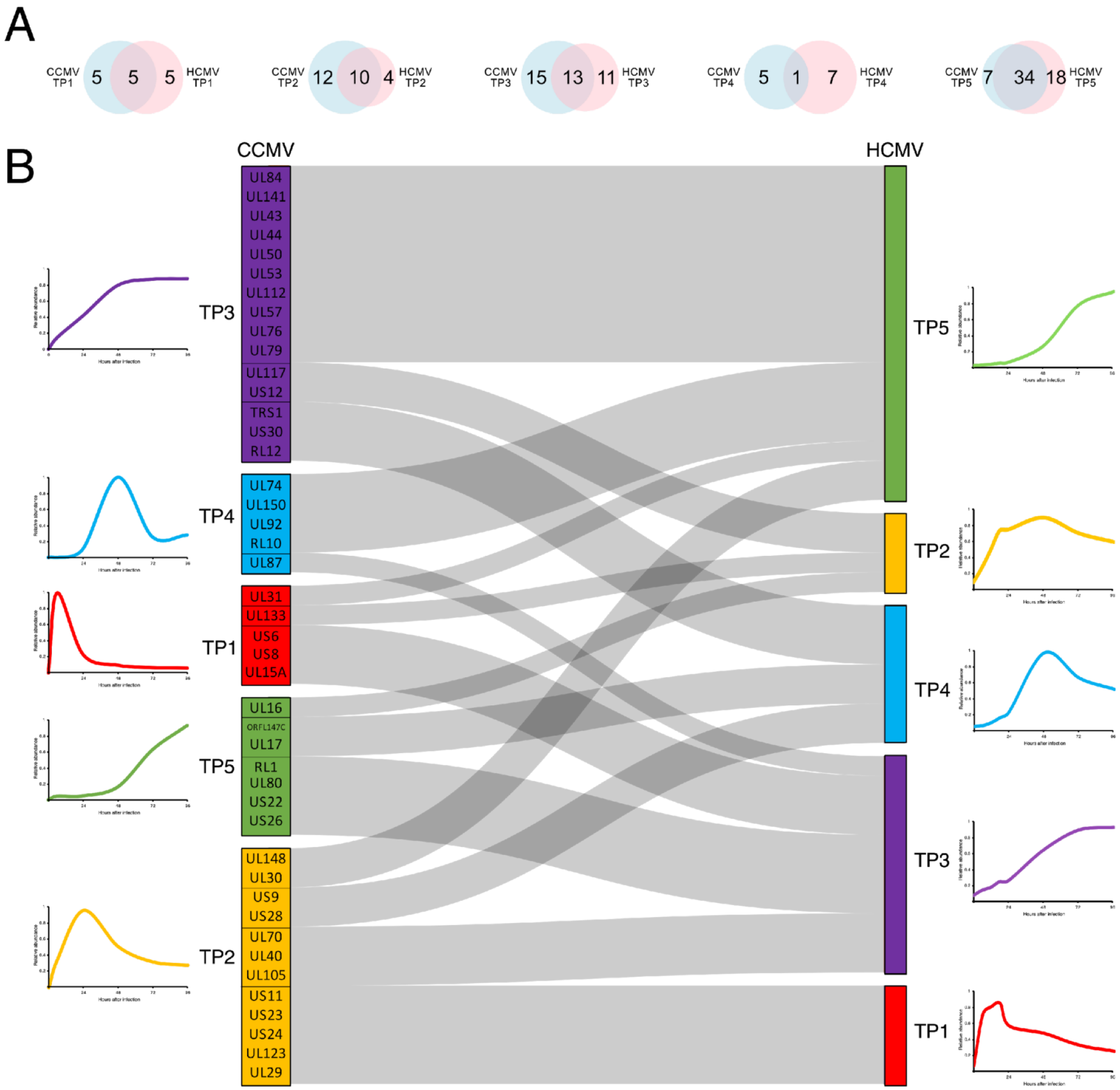


**FIG S10** TP class comparison between HCMV and CCMV. (A) Venn diagram shows the amount of overlap of homologs in each TP class. (B) Sankey diagram shows the TP class switch of homologs from CCMV to HCMV.

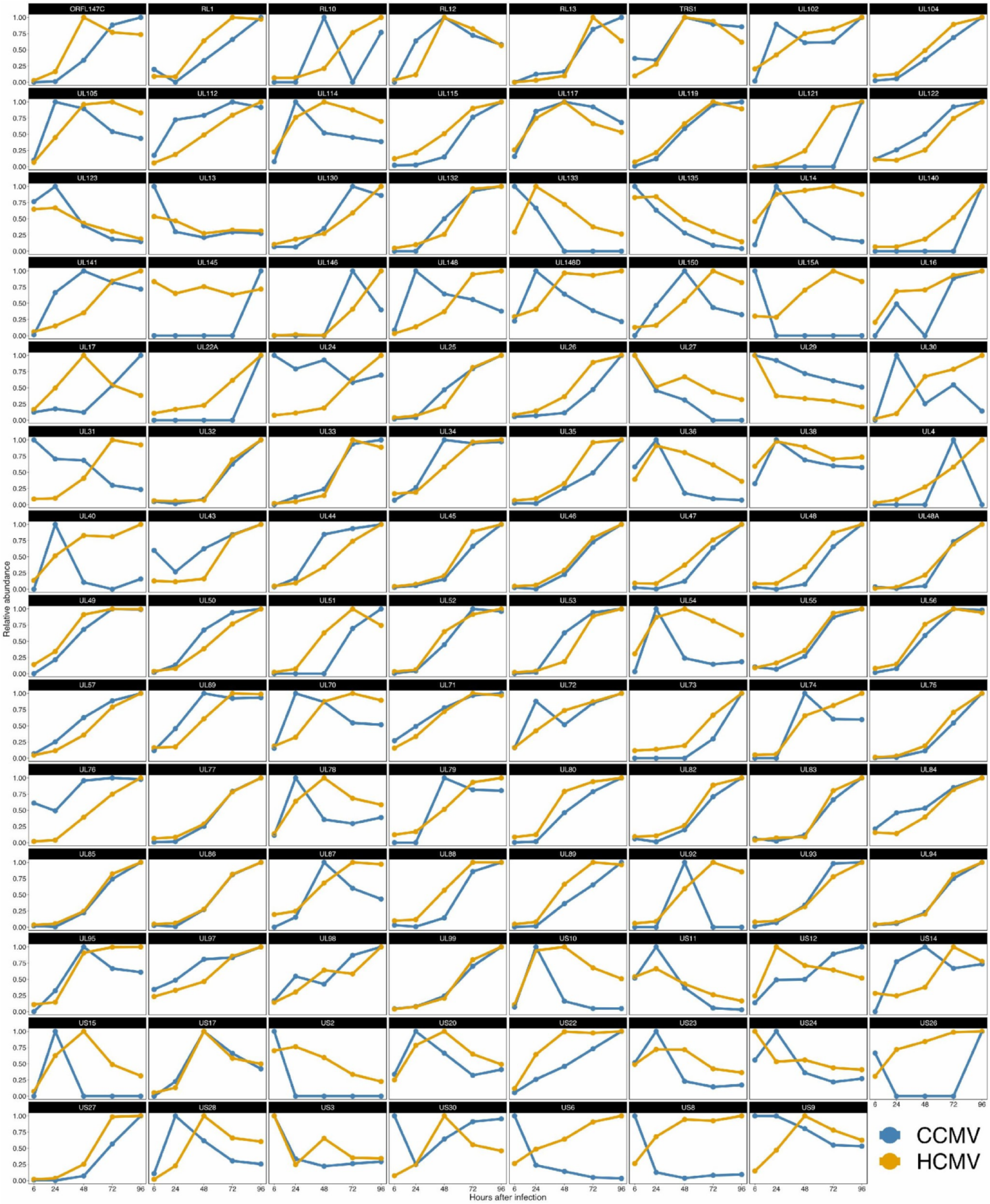


**FIG S11** Comparison of CCMV and HCMV protein expression files. HCMV data were taken from Weekes *et al*. 2014.

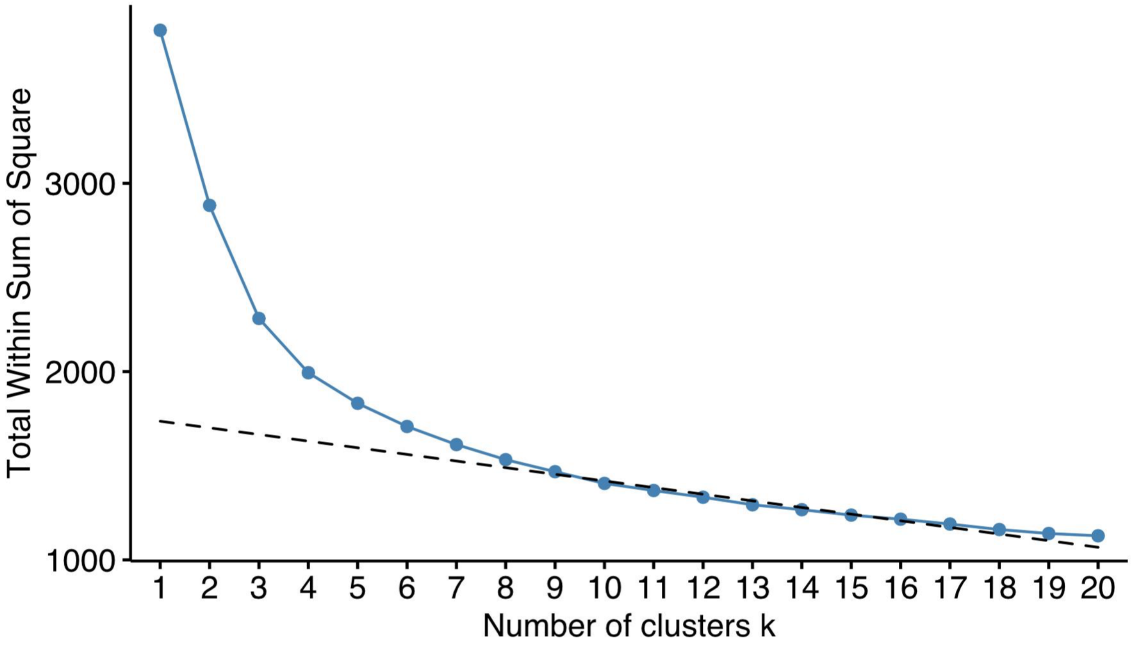


**FIG S12** Determining the number of temporal classes of host RNA/protein expression during CCMV infection. The summed distance of each gene from its cluster centroid was calculated for one to 20 clusters and plotted. The point of inflection fell at seven.

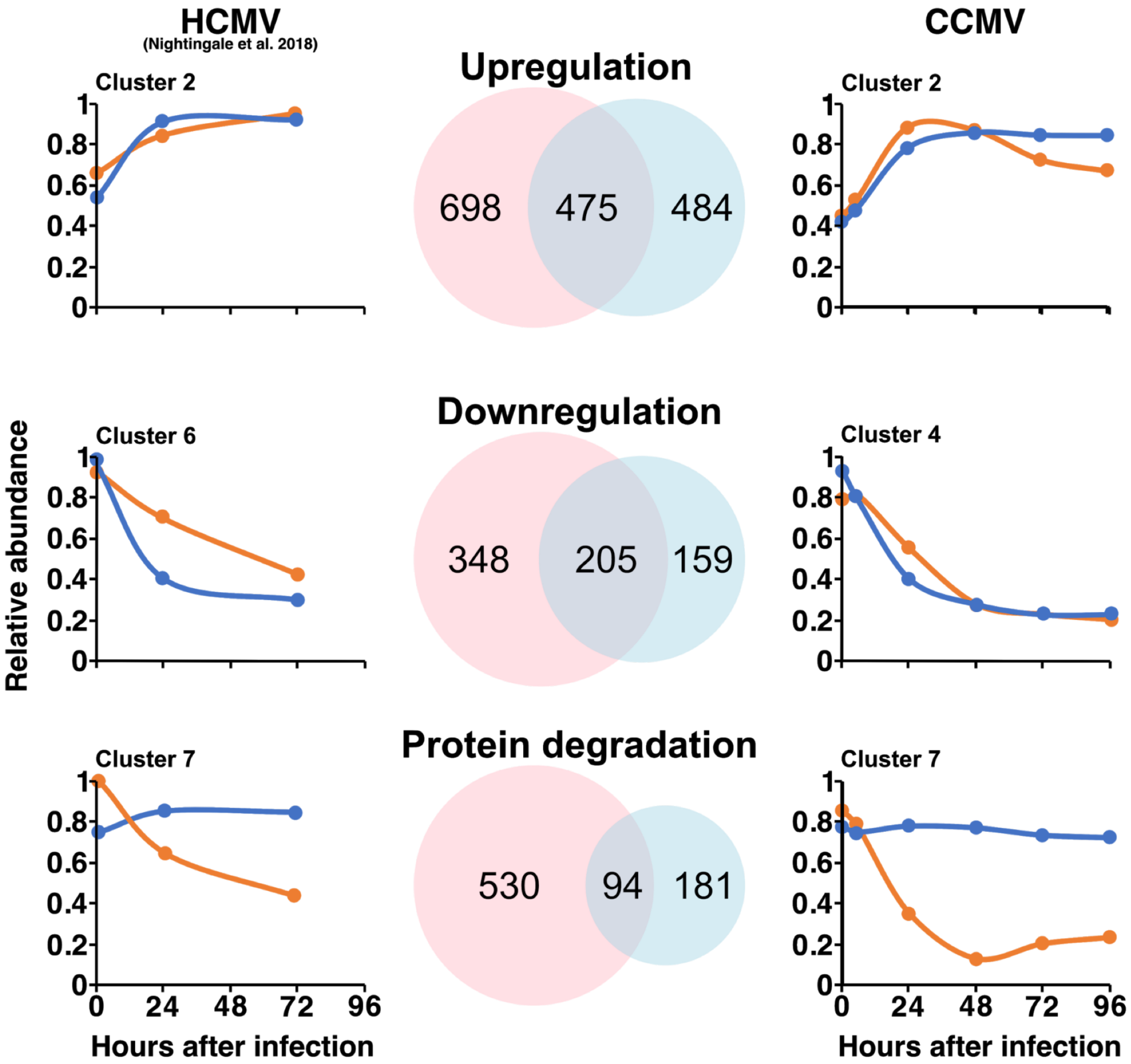


**FIG S13** Cellular factors with similar expression profiles during HCMV and CCMV infection. Selected clusters with kinetics that indicate biological relevant modes of regulation (Transcriptional up- and downregulation, protein degradation) were compared between CCMV and HCMV data sets (Nightingale et al. 2018). Left and right panels show the average cluster profiles. Venn diagrams show the overlap of genes in paired clusters.
